# A descending pathway facilitates undulatory wave propagation in *Caenorhabditis elegans* through gap junctions

**DOI:** 10.1101/131490

**Authors:** Tianqi Xu, Jing Huo, Shuai Shao, Michelle Po, Taizo Kawano, Yangning Lu, Min Wu, Mei Zhen, Quan Wen

## Abstract

Descending signals from the brain play critical roles in controlling and modulating locomotion kinematics. In the *Caenorhabditis elegans* nervous system, descending AVB premotor interneurons exclusively form gap junctions with B-type motor neurons that drive forward locomotion. We combined genetic analysis, optogenetic manipulation, and computational modeling to elucidate the function of AVB-B gap junctions during forward locomotion. First, we found that some B-type motor neurons generated intrinsic rhythmic activity, constituting distributed central pattern generators. Second, AVB premotor interneurons drove bifurcation of B-type motor neuron dynamics, triggering their transition from stationary to oscillatory activity. Third, proprioceptive couplings between neighboring B-type motor neurons entrained the frequency of body oscillators, forcing coherent propagation of bending waves. Despite substantial anatomical differences between the worm motor circuit and those in higher model organisms, we uncovered converging principles that govern coordinated locomotion.

**Significance Statement:** A deep understanding of the neural basis of motor behavior must integrate neuromuscular dynamics, mechanosensory feedback, as well as global command signals, to predict behavioral dynamics. Here, we report on an integrative approach to defining the circuit logic underlying coordinated locomotion in *C. elegans.* Our combined experimental and computational analysis revealed that (1) motor neurons in *C. elegans* could function as intrinsic oscillators; (2) Descending inputs and proprioceptive couplings work synergistically to facilitate the sequential activation of motor neuron activities, allowing bending waves to propagate efficiently along the body. Our work thus represents a key step towards an integrative view of animal locomotion.

## Introduction

Locomotion requires coordinated rhythmic motor activity. In vertebrate and invertebrate motor systems, oscillatory signals that innervate muscles are generated by dedicated neurons and neural circuits with intrinsic rhythmic properties, called central pattern generators (CPGs) (1-6). Movements, which constitute muscle activities of defined spatial pattern and temporal sequence, are driven by CPG modules distributed at different body segments. Coordination of rhythmic movements requires coupling mechanisms, by which the rhythm of multiple CPGs becomes frequency-and phase-locked (5, 7).

Although the activity of CPGs is intrinsic and independent of sensory inputs, proprioceptive or mechanosensitive feedback reshapes the motor dynamics during movement (8). In lamprey and leech, for example, activation of specialized proprioceptive neurons, either by current injection or externally imposed body bending, entrains the motor CPG activity (9, 10). In limbed animals, rapid phasic feedback from mechanoreceptors tunes the rhythmic firing of motor neurons during step cycles (11-13). Mice lacking Piezo2, a mechanically activated channel expressed in proprioceptive neurons, exhibit severely uncoordinated body movement (14).

Descending signals, typically the projections of glutamatergic neurons from the brain to the spinal or nerve cords, regulate both CPG activity and motor patterns (15). In fish and rodent, descending reticulospinal neurons, arising from the brainstem and innervating excitatory spinal interneurons, can initiate locomotion and modulate speed (16-19); whereas those that activate inhibitory spinal interneurons can terminate locomotion (20). Descending interneurons that directly target the forelimb motor neurons are critical for skillful motor behaviors (21). In leech, activation of a descending command interneuron, cell 204, can initiate and maintain swimming behavior (7, 22).

A deep understanding of the motor circuit must integrate local CPG elements and sensory feedback, as well as global command signals, to predict behavioral dynamics from interaction of individual circuit components (23, 24). With a connectome at synaptic resolution and fully identified cell-types (25, 26), *Caenorhabditis elegans* offers an opportunity to obtain a complete, system-level understanding of a locomotor circuit.

The *C. elegans* ventral nerve cord consists of a network of excitatory cholinergic motor neurons, the A-and B-type that drive backward and forward movement, respectively, and the inhibitory motor neurons (D-type) that are innervated by the excitatory motor neurons (27). The AVB premotor interneurons, which receive numerous inputs from sensory and other interneurons, extend long processes along the entire ventral nerve cord, and exclusively form gap junctions with the B-type motor neurons (26, 28, 29) (Fig. 1*A*). The functions of AVB premotor interneurons and their electrical synapses with B-type motor neurons, however, are poorly understood.

**Fig. 1.**
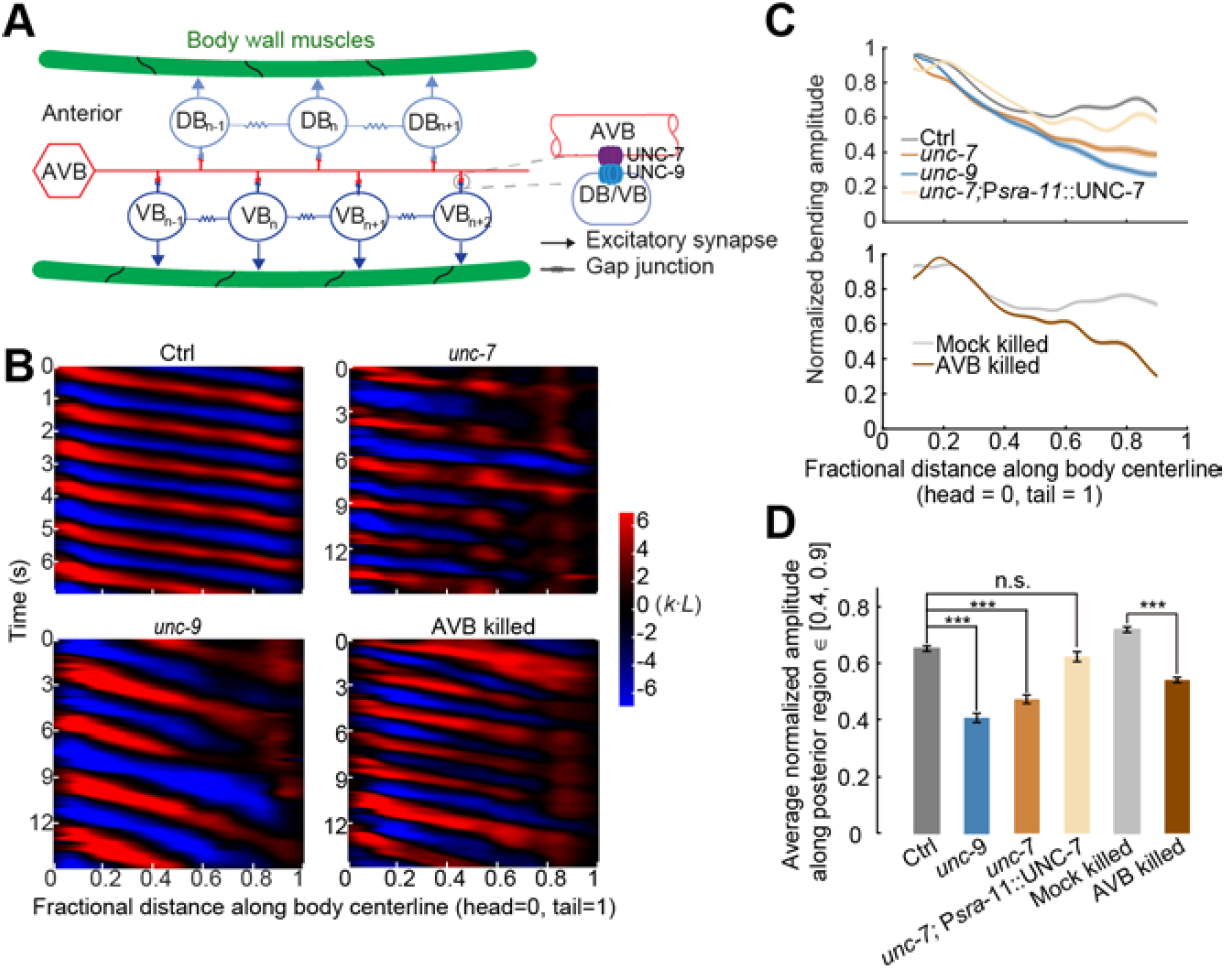
AVB-B gap junctions facilitate undulatory wave propagation during forward locomotion. (*A*) Schematics of gap junction couplings in the *C. elegans* forward motor circuit. The AVB premotor interneurons form gap junctions with all B-type motor neurons, which drive dorsal (DB) and ventral (VB) muscle contraction. AVB-B gap junctions require the expression of innexin UNC-7 in AVB interneurons and UNC-9 in B-type motor neurons. Neighboring B-type neurons are also gap junction coupled. Other types of motor neurons have been omitted from the diagram for simplicity. (*B*) Representative curvature kymographs of control (N2; P*unc-4*::TWK-18(gf)), *unc-7* (*unc-7*(*hp121*); P*unc-4*::TWK-18(gf)), *unc-9* (*unc-9*(*fc16*); P*unc-4*::TWK-18(gf)) animals, and AVB-ablated (P*lgc-55*::miniSOG; P*unc-*4::TWK-18(gf)) and their control (Mock-ablated) animals. Worms swam in viscous solutions (∽ 1 Pa·s viscosity). Body curvature was defined as a non-dimensional unit *k*·*L*, where *L* is the body length. (*C*) Quantification of the bending amplitude along the body for strains described above. Shaded regions represent s.e.m. Amplitude was normalized against the maximum curvature along the body. (*D*) Averaged bending amplitude in the posterior body ∈ [40, 90] for strains described above. *** *p* < 0.0001, two-sample *t*-test with Bonferroni correction, n.s. *p* = 0.16. Error bars represent s.e.m. Ctrl, *n* = 105 measurements, 13 worms; *unc-7* mutant, *n* = 149 measurements, 22 worms; *unc-9* mutant, *n* = 93 measurements, 17 worms; AVB-ablated, *n* = 80 measurements, 13 worms; Mock (same genotype without ablation), *n* = 96 measurements, 12 worms.

Here, we took an integrative approach to defining the circuit logic underlying coordinated locomotion in *C. elegans*, with an emphasis on how descending signals modulate the dynamics of the motor circuit. Whereas descending inputs in other systems directly drive or modulate the activity of locomotor CPGs (16-18), it remains unclear whether similar CPG elements exist in *C. elegans* (30). Combining molecular genetics and optical manipulation of neuronal activity in freely behaving *C. elegans*, we found that mid-body B-type motor neurons could function as oscillators. To generate coherent body undulation, the intrinsic activity in B-type motor neurons was entrained by proprioceptive feedback. To promote forward movement, AVB-B gap junctions drove the bifurcation of B-type neuron dynamics from stationary to rhythmic activity.

Previously, we demonstrated the proprioceptive properties in B-type motor neurons: directional proprioceptive couplings between B-type motor neurons are responsible for the propagation of rhythmic head bending along the body (31, 32). Our current experimental and computational analyses suggest that descending signals from the AVB premotor interneurons function synergistically with proprioceptive couplings to facilitate sequential activation of motor neuron activities, allowing body bending to propagate directionally and efficiently to drive forward locomotion.

## Results

### Gap junctions between descending AVB premotor interneurons and B-type motor neurons facilitate bending wave propagation during forward locomotion

The AVB premotor interneurons and B-type motor neurons form heterotypic gap junctions via the UNC-7 innexin (in AVB) and UNC-9 innexin (in B-type) (Fig. 1*A*) (28, 29). To determine whether AVB-B gap junctions play a role in forward locomotion, we first examined the motor behaviors of *unc-7*(*hp121*) and *unc-9*(*fc16*) null mutants immersed in viscous solution (1 Pa·s viscosity). In these gap junction-deficient mutants, the intrinsically higher levels of activity in the A-type motor neurons, which control backward locomotion, prevent animals from generating continuous forward movement (29). To reduce interference from motor activity arising from the backward motor circuit, A-type motor neurons were constitutively silenced by an active K^+^ channel [P*unc-4*::TWK-18(gf)] in examined strains and experiments (Table S1).

We visualized and quantified *C. elegans* locomotion kinematics by curvature kymographs (Fig. 1*B*). During bouts of forward movement, each body segment alternated between positive (red) and negative (blue) curvature, and the stripes of curvature propagated from the head to the tail. Coordinates along the worm body were defined as head = 0 and tail = 1 (Fig. 1*B*). In control animals (wild-type with A-type motor neurons constitutively silenced), the bending amplitude was highest near the head, then declined gradually and plateaued at the mid-body (∽ 40% of the body length, Fig. 1*C*). Both *unc-7* and *unc-9* mutants exhibited identical phenotypes: their bending amplitudes diminished monotonically towards the tail (Fig. 1*B* and *D* and Video 1).

UNC-7 and UNC-9 are involved in gap junction couplings in many neurons besides AVB and B-type neurons (28, 33). We next showed that disruption of AVB-B gap junctions was responsible for the monotonic decline of bending amplitudes in *unc-7* and *unc-9* mutants. First, when AVB premotor interneurons were optogenetically ablated using miniSOG (34) (by either P*lgc-55(B)*::miniSOG or P*sra-11*::miniSOG), we observed the same trend of bending amplitude decline towards the tail (Fig. 1*B* and *D* and Fig. S1). Second, restoring innexin UNC-7 in the AVB interneurons (*unc*-7; P*sra-11*::UNC-7) in *unc-7* mutants was sufficient to restore the plateau phase of bending amplitude (Fig. 1*C* and *D* and Video 3).

### Linear model predicts deteriorated undulatory waves in the presence of AVB-B gap junctions

We next asked why AVB-B gap junction-deficient worms could not efficiently propagate body bending towards the tail. Previously, we showed that worm undulation during forward locomotion requires local and directional proprioceptive coupling between adjacent body regions (31). Curvature change in a mid-body region, induced by a pneumatic microfluidic device, activates bend-sensitive B-type neurons and defines the curvature of the posterior neighbor (31). Consistently, when we trapped the middle body region of a wild-type worm in a static channel with defined curvature, the unrestrained posterior body region exhibited bending curvature in the same direction as that imposed by the microfluidic channel (Fig. 2*A* and *B* and (31)).

**Fig. 2.**
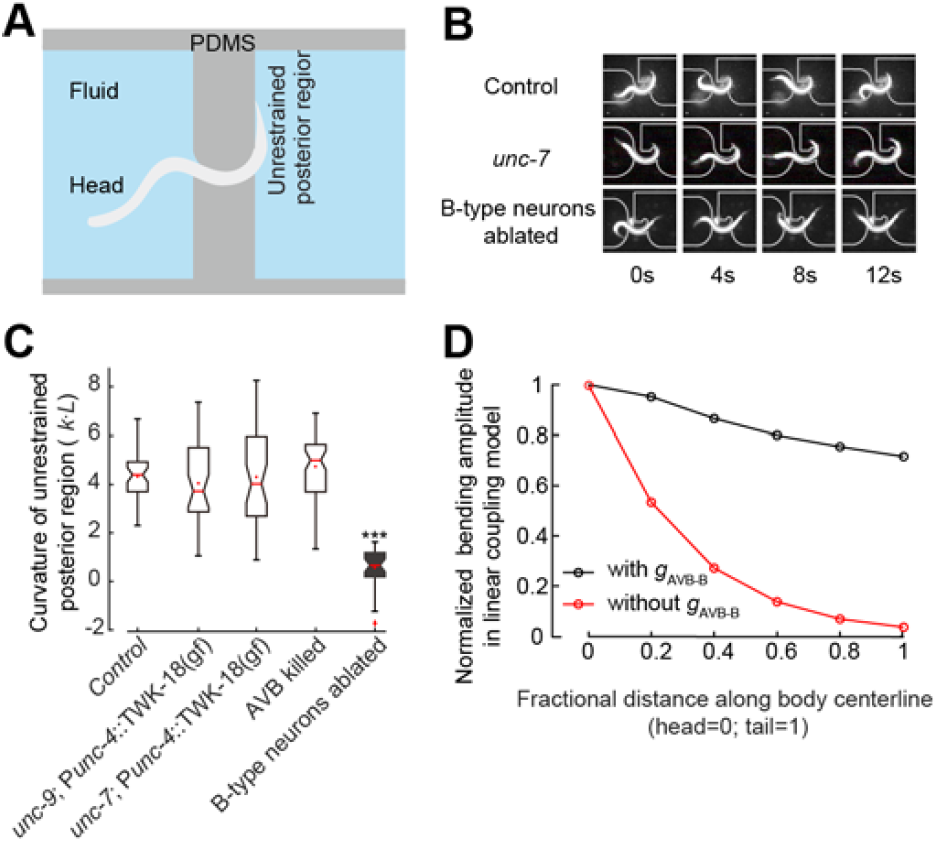
UNC-7-and UNC-9-dependent gap junctions are not required for proprioceptive couplings in the forward motor circuit. (*A*) Schematics of the microfluidic device for constraining a body segment with defined curvature. (*B*) Time-lapse video images of control (upper panel), *unc-7* (middle panel), and B-type neuron-ablated (lower panel) worms trapped in the microfluidic device. In both control and *unc-7* mutant animals, the posterior unrestrained body region followed the curvature of the channel. However, this was not the case when B-type motor neurons were optogenetically ablated. (*C*) In AVB-B gap junction-deficient mutants, or AVB-ablated worms, the posterior unrestrained body region also followed the curvature of the channel. Boxes indicate Q1 to Q3, error bars represent Q3 + 1.5IQR respectively, notches indicate 95% confidence interval. *** *p* < 0.0001, compared with other strains, by Mann-Whitney U test. Control; (N2; P*unc-4*::TWK-18(*gf*)), *n* = 40 measurements, 8 worms; *unc-7* (*unc-7;* P*unc-4*::TWK-18(gf)), *n* = 41 measurements, 12 worms; *unc-9* (*unc-9;* P*unc-4*::TWK-18(gf)), *n* = 38 measurements, 9 worms; AVB-ablated (P*lgc-55(B)*::miniSOG), *n* = 60 measurements, 10 worms; B-ablated (P*acr-5*::miniSOG), *n* = 33 measurements, 9 worms. (*D*) In the linear model, incorporating the AVB-B gap junction inputs further deteriorated bending wave propagation.

To determine whether the proprioceptive coupling is UNC-7-and UNC-9-dependent, we examined *unc-7* and *unc-9* mutants constrained by microfluidic channels. Consistent with a previous finding (31) and observation from control animals, their posterior body regions still followed the curvature imposed on the anterior body regions (Fig. 2*B and C* and Fig. S2). Therefore, proprioceptive coupling remains largely intact in the absence of UNC-7-or UNC-9-dependent gap junctions.

We next took a theoretic approach to probing how AVB-B gap junctions might affect bending wave propagation. We first adopted a linear model (31), where we asserted that the undulatory wave started with rhythmic dorsal / ventral bends near the head. Directional proprioceptive coupling between adjacent body regions is fully described by a set of first order differential equations:

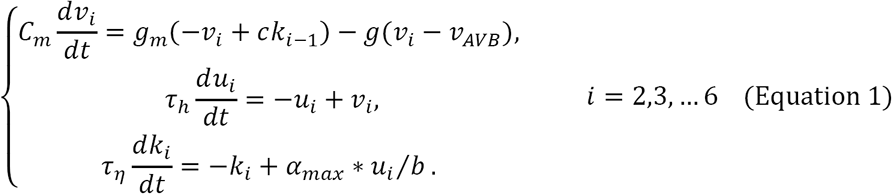

We divided the whole worm into several segments, with subscript *i* as the segment number. In the first equation, *v* describes relative motor neuron activity: when dorsal B-type neurons have a higher activity than ventral ones, *v* > 0, and vice versa. *k*_*t*_ is the curvature of a given segment, and *ck*_*t*-1_ describes the proprioceptive signal from the neighboring anterior body segment, where *c* > 0 is the coupling strength. *C*_*m*_ is the membrane capacitance and *g*_*m*_ is the leaky conductance. The term -*g*(*v*_**i**_-*v*_**AVB**_) describes the contribution from AVB-B electrical coupling, where *g* is the gap junction conductance. Here we set AVB membrane potential at a constant depolarized value, which is consistent with experimental observations that AVB interneurons exhibit elevated yet non-oscillatory calcium activity during forward locomotion (29, 35, 36). In Equation 1, for simplicity, we set *v*_*AVB*_ = 0 to eliminate the bias of *v*. A more detailed model that treated dorsal and ventral motor neuron activities separately is described in the Supporting Information.

The second equation relates motor neuron activity to muscle torque: *u* is a dimensionless variable and α_**max**_ ∗ *u* is the maximum torque that the muscle cells can generate. The last equation describes how the segment curvature changes with muscle torque: *b* is the bending modulus of the body and τ _*η*_ is a time constant (31). Published work (31, 37, 38) has established that when *C. elegans* swam in viscous solution (our experimental condition), the biomechanics of the worm body can be modeled as elastic rod bent in viscous medium at low Reynolds number, and the inertia can be ignored. If we consider sinusoidal dynamics of curvature and torque, the time constant τ_*η*_ has the following expression (31, 37):

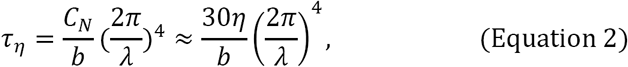

where *C*_**N**_ is the drag coefficient perpendicular to the worm body, a parameter that is proportional to the viscosity of medium η, and λ is the spatial wavelength of undulation.

Simulation of the linear model predicted an exponential decay of bending amplitude towards the tail (Fig. 2*D*). By considering a continuous form of the local coupling (see Supporting Information), we solved the linear model analytically and identified a self-consistent solution for the spatiotemporal profile of the worm body curvature:

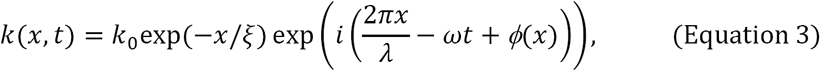

where *ω* describe the angular frequency; *φ*(*x*) is a phase lag term, and ξ is the decay length constant, which, to the leading order, is given by:

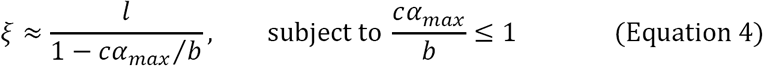

where *l* is the spatial scale for local proprioceptive coupling (31). ξ → + ∞ leads to identical bending amplitude along the body. Incorporation of AVB-B electrical coupling effectively changes *c* to *cg*_**m**_ / (*g*_**m**_ + *g*). This prefactor further reduces the length constant. In a linear coupling model where B-type motor neurons are passive recipients of proprioceptive inputs, opposite to our experimental findings, gap junctions between AVB and B-type neurons would deteriorate the propagation of bending waves (Fig. 2*D*).

### B-type motor neurons are nonlinear functional units

The discrepancy between the theoretical model and experimental observations indicates that some basic assumption in the linear model must be modified. We hypothesized that proprioceptive signals would couple nonlinear functional units within adjacent regions. Intrinsic oscillation is a common form of nonlinearity. We thus tested whether CPGs exist along the worm nerve cord.

Our linear coupling model predicted that abolishing head bending activity would abolish the undulatory wave; a coupled-CPG model (39), however, predicts the opposite. To distinguish these possibilities, we performed spatially selective optogenetic inhibition of B-type motor neurons in a defined anterior body region (0.1–0.3 along worm coordinates) of a freely swimming animal (P*acr-5*::Arch) (Fig. 3*A–B* and Video 2). Upon green light illumination, we effectively abolished bending near the head region (Fig. 3*B*, curvature kymograph). In the absence of local proprioceptive signal from adjacent anterior body region, the mid-body (∽ 50% of the body length), however, continued to generate rhythmic bending with increased undulation frequency and reduced amplitude (Fig. 3*B* and 3*E*, Video 2). When a worm swam in 1 Pa·s viscous solution, the undulation frequency was ∽ 0.8 Hz (Fig. 3*C*); when anterior bending activity was optogenetically inhibited, the mid-body undulation frequency doubled (∽ 1.6 Hz, Fig. 3*C*). We observed a similar phenomenon when anterior undulation was abolished optogenetically, either by inhibiting both A-type and B-type motor neurons (P*acr-2*::Arch, Fig. S3*A*), or by directly inhibiting anterior muscle cells (P*myo-3*::NpHR, Fig. S3*A*).

**Fig. 3.**
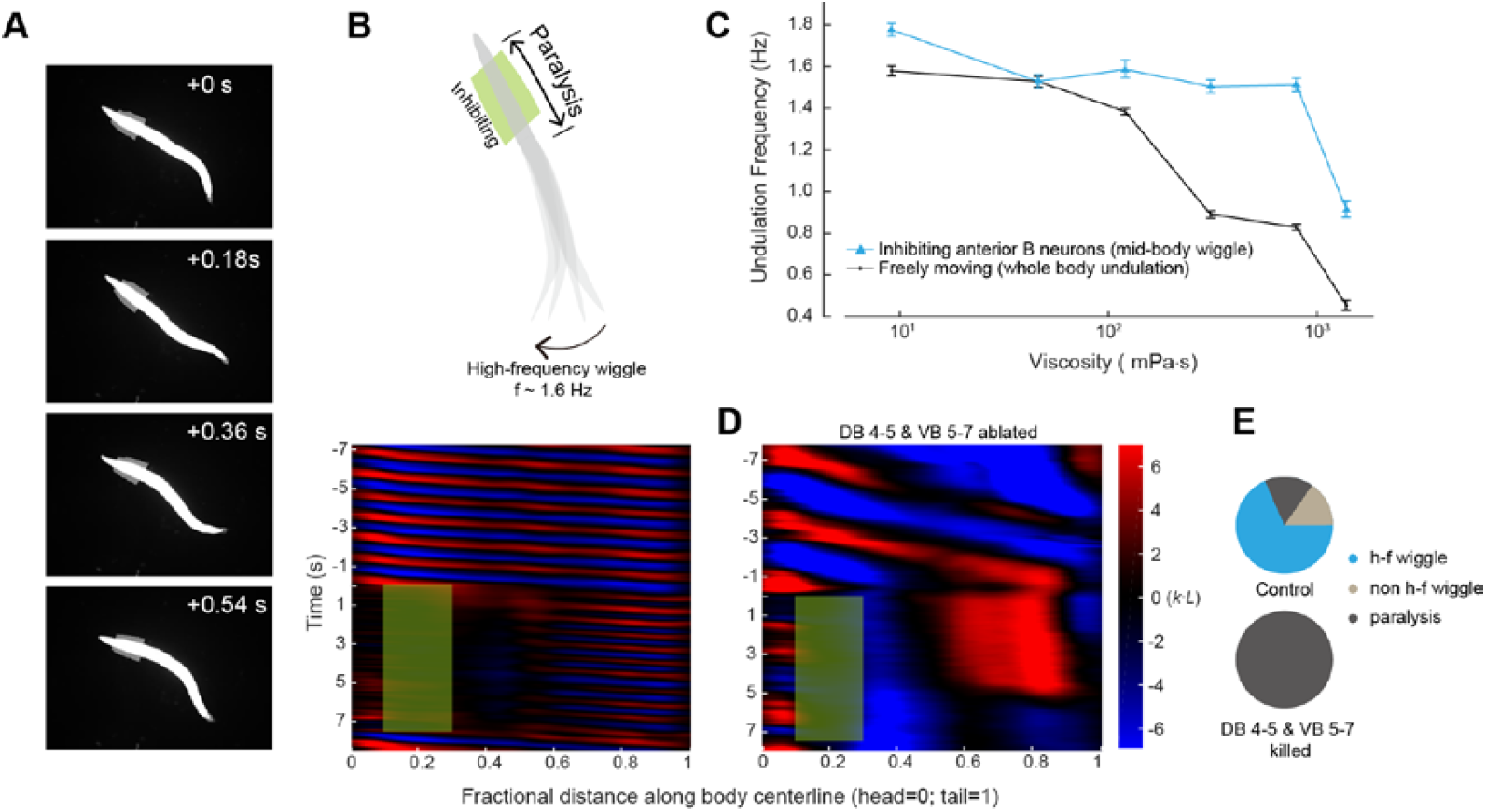
Mid-body B-type motor neurons generate rhythmic activity independent of proprioceptive coupling. (*A*) Time-lapse video images from a recording when B-type motor neurons in an anterior body region (10–30% along the worm body) were optogenetically inhibited. (*B*) Upper panel, schematic illustrates the effect of spatially selective inhibition of B-type motor neurons. Optogenetic inhibition of anterior B-type motor neurons induced high-frequency undulation in the posterior region; lower panel, representative curvature kymograph. Green shaded region shows the selected spatiotemporal region for optogenetic inhibition. (*C*) *C. elegans* undulation frequency at different viscosity. Black line is undulation frequency of control animals; blue line is mid-body undulation frequency when anterior bending activity was abolished. Error bars are s.e.m.; *n* ≥ 8 worms for each data point. (*D*) Representative curvature kymograph during optogenetic inhibition of anterior B-type motor neurons, with and without mid-body B-type neurons (DB 4–5 & VB 5–7). (*E*) Pie chart summarizes the percentage of locomotor states when anterior bending activity was abolished. h-f wiggle: mid-body undulation frequency was higher than that before anterior bending activity was abolished; non-h-f wiggle: mid-body undulation frequency was equal to or less than that before anterior bending activity was abolished; paralysis: no waves emerged in the mid-body. Control (P*acr-5*::Arch), *n* = 241 measurements, 20 worms; Mid-body B-type neuron-ablated worms (P*acr-5*::Arch; P*acr-5*::miniSOG), *n* = 77 measurements, 11 worms.

These observations favor the possibility that at least some B-type motor neurons are intrinsic oscillators. Whereas the normal undulation frequency decreased in more viscous solution (Fig. 3*C* and (37)), the mid-body undulation frequency remained fixed across a wide range of viscosities when anterior bending activity was abolished (Fig. 3*C* and Fig. S3*B*). Therefore, the intrinsic frequency of the mid-body oscillators, which could be entrained by head bending, appeared to be independent of the mechanical load of the environment.

To determine whether the B-type motor neurons were responsible for generating intrinsic mid-body oscillation, we performed systematic ablation of B-type neurons (P*acr-5*::miniSOG) in the ventral nerve cord. Ablating a cluster of mid-body B-type motor neurons (DB4-5 and VB5-7) abolished high-frequency oscillation when anterior bending activity was abolished (Fig. 3*D* and *E*). When only a subset of neurons in this cluster was ablated, the high-frequency mid-body undulation could still be induced (Fig. S3*A*). In these animals, we occasionally observed independent bending waves generated by the head and posterior body regions, respectively (Fig. S3*C*). When local inputs to B-type neurons were blocked, either by silencing A-type motor neurons via a gain-of-function potassium channel (P*unc*-4::TWK-18, Figure 4B control) or by eliminating inhibitory neurotransmitter release (Fig. S3A), intrinsic mid-body undulation persisted. These data, together with the observations that worm mid-body oscillated in the absence of local proprioceptive signal arising from adjacent anterior region (Fig. 3B and Fig. S3A), strongly suggest that B-type motor neurons do not passively respond to proprioceptive signals. They instead function as nonlinear units that resemble distributed CPG modules along the body.

**Fig. 4.**
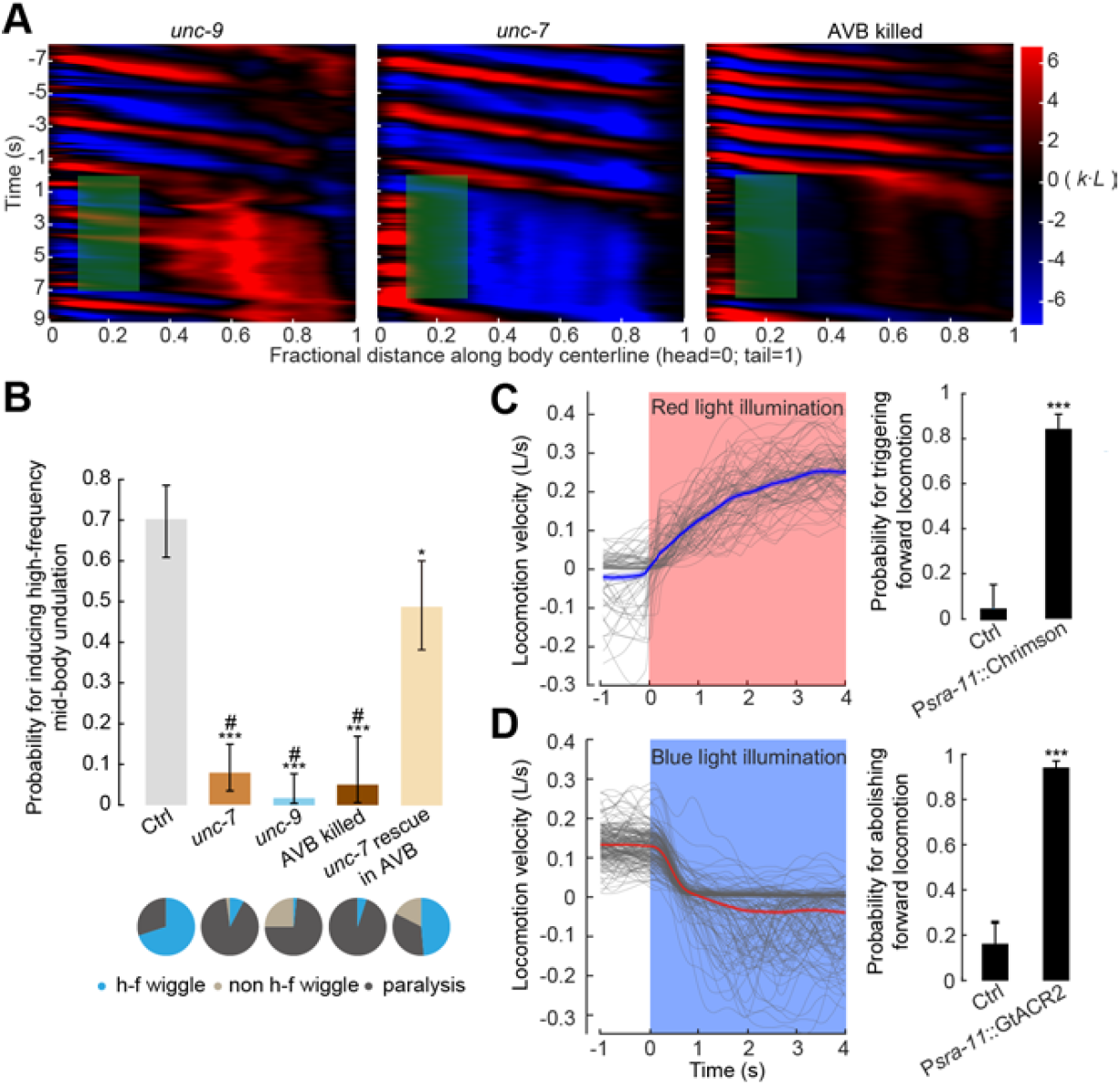
AVB-B electrical coupling drives the bifurcation in the B-type neuron dynamics. (*A*) Representative curvature kymographs in AVB-B gap junction-deficient or AVB-ablated worms when anterior bending activity was abolished. (*B*) Probability for inducing high-frequency mid-body undulation when anterior bending activity was abolished. Pie chart summarizes the percentage of worm locomotor states. Error bars indicate 95% binomial proportion confidence interval. *** *p* < 0.0001, * *p* = 0.002, # *p* < 0.0001 (* compared with Ctrl, # compared with UNC-7 rescued in AVB.) Chi-square test. Strains for comparisons are: control [P*acr-5*::Arch; P*unc-4*::TWK-18(gf)], *n* = 113 measurements, 13 worms; *unc-7* [*unc-7*(*hp121*); P*acr-5*::Arch; P*unc-4*::TWK-18(gf)] *n* = 102 measurements, 21 worms; *unc-9* [*unc-9*(*fc16*); P*acr-5*::Arch; P*unc-4*::TWK-18(gf)] *n* = 116 measurements, 26 worms; AVB-ablated [P*lgc*-*55(B)*::miniSOG, P*acr-5*::Arch] *n* = 40 measurements, 11 worms; UNC-7 rescued in AVB [*unc-7*(*hp121*); P*acr-5*::Arch; P*unc-4*::TWK-18(gf); P*sra-11*::UNC-7] *n* = 86 measurements, 12 worms. (*C*) Optogenetic activation of AVB premotor interneurons could trigger forward locomotion. Left, optogenetic activation (P*sra*-11::Chrimson, *n* = 133 measurements, 11 worms), showing locomotion velocity with time, red light (635 nm, 12 mW / mm^2^) was turned on at *t* = 0. Gray traces represent individual trials, and shaded region represents s.e.m. Right, probability for triggering forward locomotion. (D) Optogenetic inhibition of AVB premotor interneurons could abolish forward locomotion. Left, optogenetic inhibition (P*sra*-11::GtACR2, *n* = 187 measurements, 21 worms), showing locomotion velocity with time, blue light (473 nm, 0.76 mW / mm^2^) was turned on at *t* = 0. Gray traces represent individual trials, and shaded region represents s.e.m. Right, probability for abolishing forward locomotion. Ctrl represents worms without feeding all-*trans* retinal. Error bars indicate 95% binomial proportion confidence interval, \*\*\**p* < 0.0001, Chi-square test.

### AVB-B gap junctions drive the bifurcation of B-type motor neuron dynamics

We next asked whether AVB-B gap junction inputs played a role in high-frequency mid-body undulation when anterior bending activity was abolished. Indeed, in AVB-B gap junction-deficient mutants (*unc-7* and *unc-9*), as well as AVB-ablated animals, optogenetic inhibition of anterior bending no longer induced mid-body undulation (Fig. 4*A* and *B*). Instead, optogenetic inhibition of either anterior B-type motor neurons or anterior body wall muscles led to paralysis of the whole-body (Fig. 4*A* and *B* and Fig. S4). Restoring UNC-7 innexin expression in AVB interneurons (*unc*-7; P*sra-11*::UNC-7; P*unc-4*:TWK-18(gf)) significantly increased the efficacy for inducing high frequency mid-body undulation (Fig. 4*B* and Video 3). These results suggest that AVB interneurons and their gap junction couplings with B-type motor neurons help induce mid-body high frequency undulation. Consistently, optogenetic activation of the AVB interneurons (*Psra-11*::Chrimson) reliably triggered forward locomotion from either the pause state or the backward movement (Fig. 4*C*, Video 7); whereas optogenetic inhibition of the AVB interneurons (P*sra-11*::GtACR2) (40) reliably abolished forward locomotion (Fig. 4D, Video 8).

Given that AVB interneuron activities are *non-oscillatory* (29, 35, 36), our data thus suggest that AVB-B gap junctions drive a bifurcation of B-type neuron dynamics, leading to a transition from the stationary to oscillatory state. In the presence of AVB-B gap junctions, time-varying proprioceptive signals from the anterior body may easily induce changes in motor neuron membrane potential, thus facilitating bending wave propagation.

### Nonlinear model recapitulates AVB-B gap junction-dependent bifurcation of B-type motor neuron dynamics

We next sought theoretical verification of our functional model. For simplicity, we developed a phenomenological model based on active Ca^2+^ and K^+^ conductance (41), in which the dynamics of dorsal and ventral motor neurons within a given body segment are governed by the following equations:

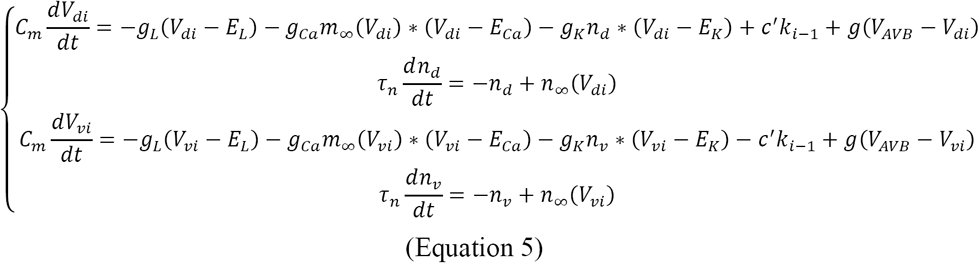

In Equation 5, *V* is motor neuron membrane potential; subscripts *d and v* denote dorsal and ventral sides, respectively; *m* and *n* are voltage-dependent Ca^2+^ and K^+^ activation variables, respectively. Parameter values in this equation were chosen (Supporting Information) to accommodate the following experimental findings. First, when the head and body undulations were decoupled (*c*'*k*_*1*_ ≈ 0), gap junctions between AVB interneurons and B-type motor neurons induced high frequency motor neuron oscillation (Fig. 5*A*). Second, proprioceptive signals arising from head bending entrained posterior motor neuron oscillation (Fig. 5*A*). The nullcline analysis, which allows for visualization of the dynamical variables on a vector field, provided useful information on the transition from the resting state to the limit cycle in the presence of AVB-B gap junctions (Fig. 5*B*). Furthermore, simulation recapitulated the experimental finding that AVB-B gap junctions helped equalize bending amplitude along the body (Fig. 5*C*).

**Fig. 5.**
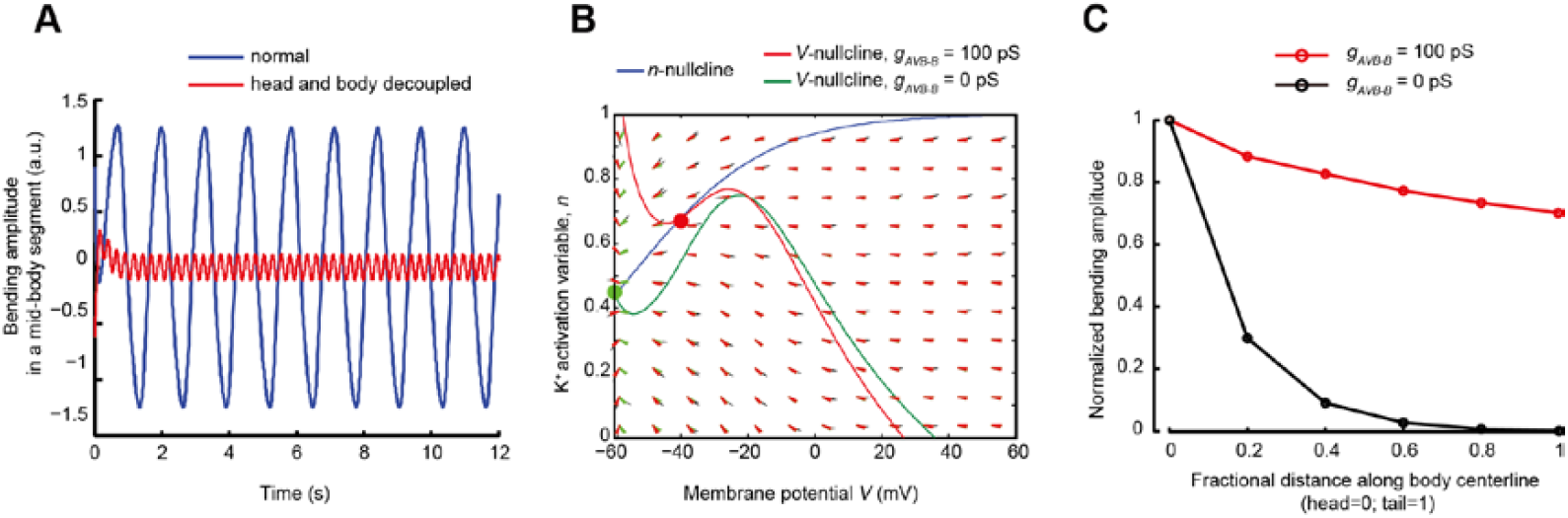
Simulation of a nonlinear model. (*A*) Bending activity of a mid-body segment. Blue, bending amplitude during normal locomotion; red, when the head and body undulation was decoupled, oscillation with higher frequency and lower amplitude emerged. (*B*) Nullcline analysis of motor neuron dynamics. The *n*-and *V*-nullclines were defined as 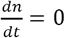, 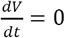, respectively. In the absence of proprioceptive coupling and AVB-B gap junctions (*g*_*AVB-B*_ = 0), the fixed point of the membrane potential dynamics (green) corresponds to a steady state; in the presence of AVB-B gap junctions, that is, *g*_*AVB-B*_ = 100 *p*s, the system undergoes bifurcation, and the new fixed point becomes unstable (red), leading to periodic oscillation. (*C*) A comparison of bending amplitude along the body, with and without AVB-B gap junctions. All parameters used in the simulation are described in the Supporting Information.

### Electrical couplings between motor neurons and AVB-B gap junctions permit rapid and reciprocal interactions between head and body motor activities

Having considered the functions of the AVB-B gap junctions, we next asked whether local gap junctions between B-type motor neurons (Fig. 1*A*), identified by anatomical wiring (25, 26), played additional roles in forward locomotion.

This question arises from our observation that inhibition of B-type motor neurons in middle or posterior body regions, unlike the anterior body region, induced rapid (within ∽ 300 ms) paralysis of the whole worm (Fig. 6*A* and *B*, Fig. S6, and Video 4). Electrical coupling is the best candidate for rapid redistribution of a hyperpolarization current among B-type motor neurons (P*acr-5*::Arch). In both *unc-7* mutants and AVB-ablated worms, whole body paralysis could still be induced upon silencing mid-body and posterior B-type motor neurons (Fig. 6*C*) using the maximum laser intensity, suggesting that the hyperpolarization signal could still be transduced in the absence of AVB-B gap junctions. In *unc-9* mutants, however, rhythmic bending in the anterior body persisted when mid-body B-type motor neurons were silenced (Fig. 6*A*, 6*C* and Video 5). Restoring UNC-9 expression in cholinergic motor neurons, including the B-and A-type motor neurons (P*acr-2*::UNC-9), restored the ability to optogenetically induce whole body paralysis (Fig. 6*C*). Therefore, electrical couplings among motor neurons contributed to the paralytic effect.

**Fig. 6.**
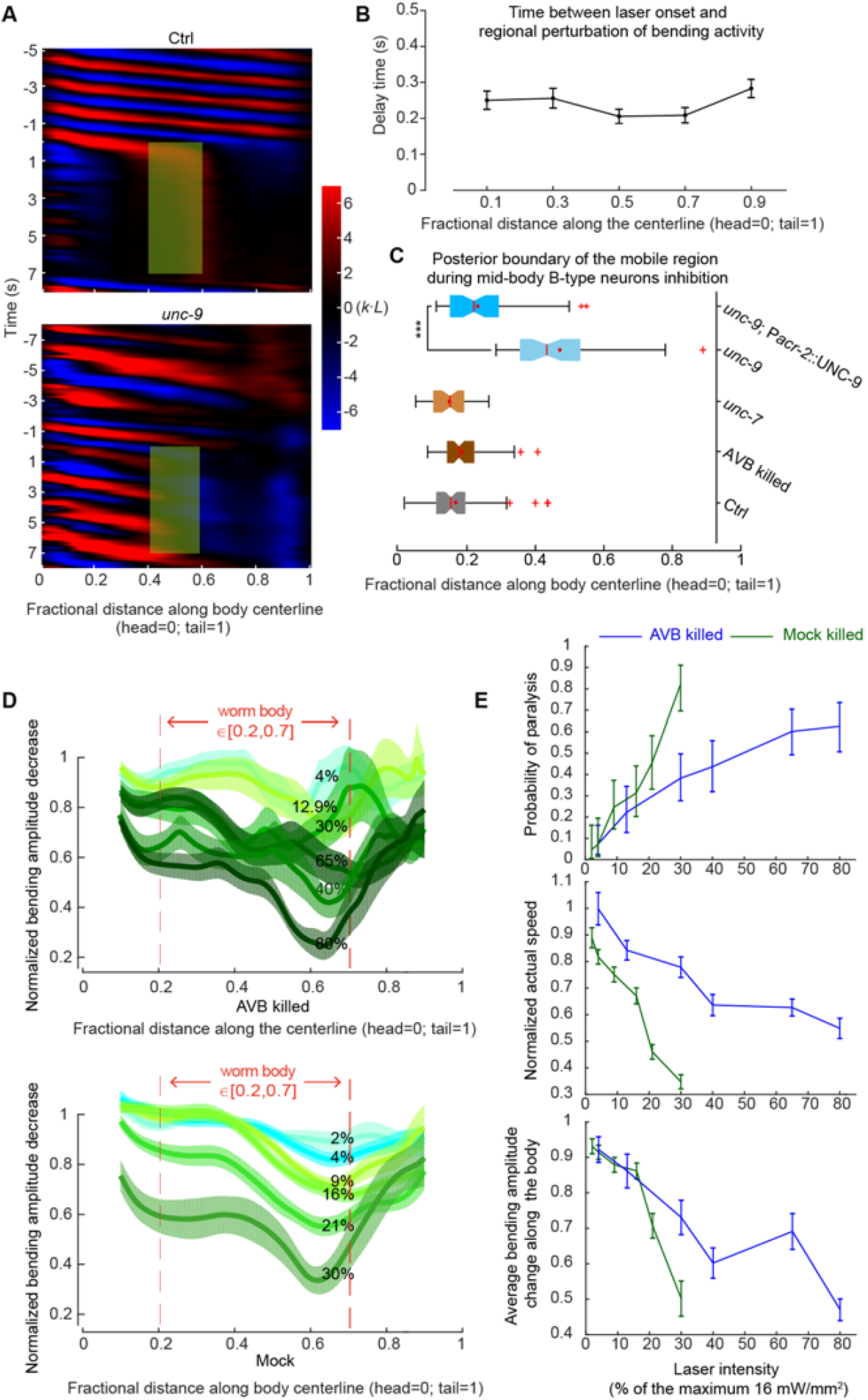
Electrical coupling among motor neurons and AVB-B gap junctions allows for rapid and reciprocal interactions between head and body motor activities. (*A*) Comparison of representative curvature kymographs of a control [P*acr-5*::Arch; P*unc-4*::TWK-18(gf)] and *unc-9* mutant [*unc-9*(*fc16*); P*acr-5*::Arch; P*unc-4*::TWK-18(gf)] worm during optogenetic inhibition of mid-body B-type motor neurons. Green shaded regions are selected regions (0.4 – 0.6 fractional distance) subjected to 0 – 7 s stimulation by the green laser. (*B*) Delay time between the onset of green laser and changes in bending. Error bars are s.e.m., also see Fig. S6. (*C*) Quantification of the whole-body paralytic effect during optogenetic inhibition of mid-body B-type motor neurons. Red crosses are outliers, red lines are medians, red dots are means, error bars represent Q1-1.5IQR and Q3+1.5IQR, respectively, and notches indicate 95% confidence interval. In *unc-9* mutants, the boundary shifted towards the mid-body, reflecting that the anterior body could still undulate. *** *p* < 0.0001, Mann-Whitney U test. Control [P*acr-5*::Arch; P*unc-4*::TWK-18(gf)], *n* = 70 measurements, 17 worms; *unc-7* [*unc-7*(*hp121*); P*acr-5*::Arch; P*unc-4*::TWK-18(gf)] *n* = 43 measurements, 11 worms; *unc-9* [*unc-9*(*fc16*); P*acr-5*::Arch; P*unc-4*::TWK-18(gf)] *n* = 59 measurements, 17 worms; AVB-ablated (P*lgc*-*55*::miniSOG, P*acr-5*::Arch) *n* = 65 measurements, 18 worms; UNC-9 rescued in the B-type and A-type motor neurons [*unc-9*(*fc16*); P*acr-5*::Arch; P*acr-2*::UNC-9] *n* = 48 measurements, 9 worms. (*D*) Laser intensity-dependent effects on the bending amplitude along the body. Error bars are s.e.m. (*E*) Laser intensity-dependent effects on efficacy for inducing whole-body paralysis, locomotion speed and mean amplitude decreases along the body [0.2, 0.7]. Error bars (speed and amplitude) are s.e.m., error bars (paralysis) are 95% binomial proportion confidence intervals. *n* ≥ 42 measurements, ≥ 6 worms for each data point. Maximum laser intensity: 16 mW / mm^2^.

The probability for optogenetically inducing whole-body paralysis, however, was dose-dependent. When the mid-body B-type motor neurons were inactivated at lower laser intensity, which likely reduced the degree of hyperpolarization, forward locomotion was not abolished. Instead, both undulation amplitude and locomotor speed were reduced (Fig. 6*D* and *E*). Interestingly, in the absence of AVB interneurons, a higher laser intensity was required to modify the locomotor kinematics and to induce whole worm paralysis (Fig. 6*D* and *E*). These results suggest a potential mechanism by which gap junctions between motor neurons and AVB-B gap junctions function together to retrogradely regulate head bending activity, and hence reconfigure the dynamics of worm body undulation.

## Discussion

Electrical synapses are prevalent in invertebrate motor circuits, as well as in developing and mature vertebrate spinal cords (42-44). The functional implications of gap junctions in the context of motor circuits and behavior can be counterintuitive, confounding, and are often underestimated (45, 46). For example, in the mature zebrafish spinal cord, electrical couplings between motor neurons and premotor interneurons point to a nimble feedback mechanism for modulating rhythmic activity (47). Experiments and modeling of crustacean stomatogastric ganglions suggest that the interplay between electrical and chemical synapses provides degenerate circuit mechanisms for switching between fast and slow oscillatory behaviors (48). In the reversal motor circuit of *C. elegans*, the electrical coupling between AVA premotor interneurons and A-type motor neurons provides state-dependent effects. At rest, they exhibit a shunting effect, by which the activity level in the backward locomotion circuit is suppressed so that a bias towards forward movement can be established (29). Upon stimulation, these gap junctions allow antidromic-rectification of A-type motor neuron activity that prolongs reversal movement (49).

In the motor circuit that drives *C. elegans* forward movement, we discovered that AVB-B electrical couplings induced intrinsic oscillations in mid-body B-type motor neurons, and facilitated undulatory wave propagation. To better understand how descending inputs drove the bifurcation of motor neuron dynamics, we developed a nonlinear dynamic model based on voltage dependent conductance. Detailed characterization of the intrinsic membrane conductance and experimental test of our model required electrophysiological recordings in motor neurons, a difficult technique that has only recently become possible (49). Intriguingly, a complementary study of the *C. elegans* reversal motor circuit found that voltage-dependent Ca^2+^ channels were required for generating intrinsic rhythmic activity in the A-type motor neurons (Shangbang Gao and Mei Zhen, personal communication, May 2017). Whether the same channels drive B-type motor neuron oscillation will be addressed in future studies.

Coordinated movement in larger animals involves networks of CPGs that are coupled through mechanosensory feedback (5). In leech, stretch-sensitive feedback can alter the intersegmental phase lags between CPGs (10, 50). During stick insect walking, proprioceptive feedback can entrain the motor patterns that drive individual joints, leading to interjoint coordination (13). Specific classes of mechanosensory neurons in fly larvae are required for the propagation of peristaltic waves (51, 52).

Here, our data indicate that distributed CPGs, operated by at least some B-type motor neurons, exist in the *C. elegans* ventral nerve cord. Because worms must adapt to extreme ranges of external mechanical loads imposed by a changing environment (37), directional proprioceptive signals, also transduced by B-type motor neurons (31), are essential for entraining the rhythm of body oscillators, and for propagating coherent bending waves from the head to the tail (Fig. 7). In a linear coupling model where motor neurons passively respond to proprioceptive inputs, bending amplitude would suffer an exponential decay towards the tail. Our experimental finding and theoretical analysis thus provide new insight into the interplay between sensory feedback and intrinsic oscillators within the motor circuit.

**Fig. 7.**
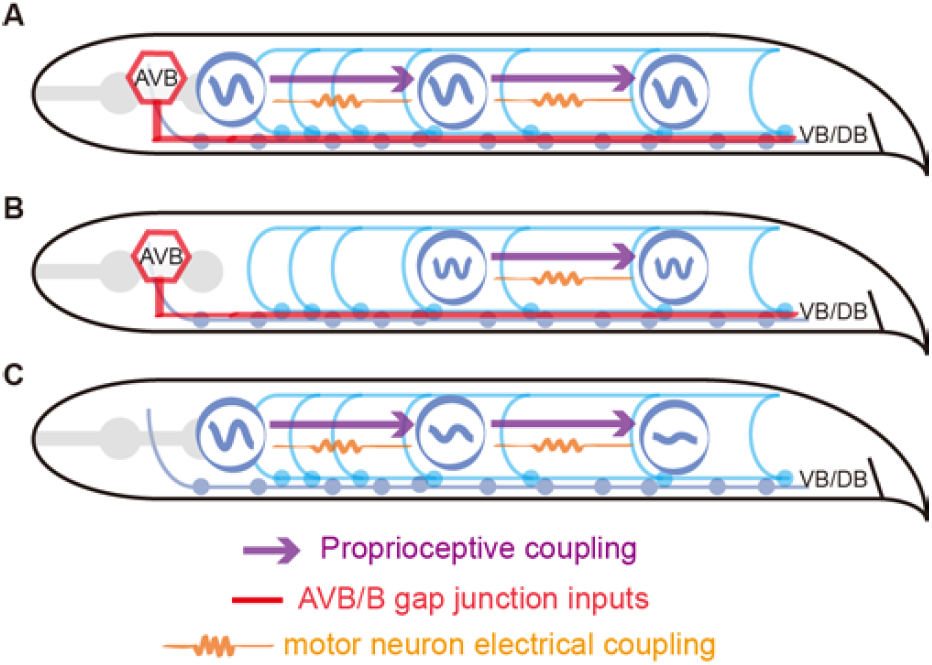
Model schematics of coordinating forward locomotion in *C. elegans*. (*A*) Proprioceptive coupling between B-type motor neurons, AVB-B gap junction coupling, and weak electrical coupling between motor neurons work synergistically to drive and propagate a coordinated undulatory wave from the head to the tail. (*B*) When a strong and time-varying proprioceptive signal from an anterior body region is absent, AVB-B gap junction coupling induces mid-body high frequency undulation. (*C*) In the absence of AVB-B gap junction inputs, proprioceptive couplings are less effective in propagating bending waves, leading to rapidly decaying bending amplitude towards the tail.

We also found a previously unknown role of electrical couplings between motor neurons, which permits rapid and reciprocal interaction between the head and body motor activities (Fig. 6 and 7). Over-expression of UNC-9 in the B-type motor neurons led to constitutive paralysis (Video 6); we suspect that strong electrical couplings between motor neurons tend to synchronize motor activity along the whole body, thus deteriorating bending wave propagation. Weak electrical couplings between motor neurons, on the other hand, might help mediate head bending activity and augment the excitability of motor neurons along the body. Direct testing of the functional contribution of local electrical couplings requires the elimination of electrical synapses between motor neurons, while sparing the AVB-B ones. Current genetic tools for manipulating gap junction expression with defined wiring specificity remain to be developed because both types of gap junctions require UNC-9 innexin.

UNC-9 hemichannels are also expressed by the body wall muscle cells (53). Could electrical couplings between muscle cells and those between motor neurons have similar functions? We (and others) found that optogenetic inhibition of mid-body muscle cells did not affect bending activity in the anterior body region, but did abolish bending activity in the posterior body region (54), consistent with our proprioceptive coupling model (Fig. 7). Furthermore, in *unc-13* mutants, where all chemical synaptic inputs from motor neurons to muscles are eliminated, targeted optogenetic activation of ventral or dorsal muscle cells induces local body bending, not bending in neighboring regions (31). These data suggest that in *C. elegans* any electrical coupling between body wall muscle cells only plays a restricted role in local bending activity.

Despite substantial anatomical differences between the worm motor circuit and those in higher organisms, we identified converging principles that govern coordinated locomotion. By integrating global descending signals, biomechanical feedback, and neuromuscular dynamics, our findings and models represent a key step towards a full system model of animal locomotion.

## Methods

### Worm strains and cultivation

Wild-type (N2), mutant, and transgenic worms were cultivated using standard methods (55). Strain information can be found in the Supporting Experimental Procedures. Transgenic worms used in all optogenetic experiments were cultivated in the dark at 20 – 25°C on NGM plates with *Escherichia coli* OP50 and all-*trans* retinal (ATR). We performed all experiments using young adult hermaphrodites.

### Microfluidic device

Custom microfluidic devices were fabricated in polydimethylsiloxane (PDMS) using soft lithography. We loaded each microfluidic channel with dextran solution [∽25% (w / w) dextran in M9 buffer (1 Pa·s viscosity)]. An individual worm was transferred into the inlet of each microfluidic channel and worm position within each channel was manually controlled by syringes connected to polyethylene tubing.

### Behavioral quantification and optogenetic manipulation

Experiments were performed on a Nikon inverted microscope (Ti-U, Japan) under 10× magnification with dark field illumination. Worms were immersed in viscous solution (∽25% (w / w) dextran in M9 buffer in most cases), sandwiched between two glass slides, and retained within the field of view of an imaging objective by a custom tracking system. Video sequences were taken by a Basler CMOS camera (aca2000-340km, Germany), and the worm body centerline was extracted in real time. We used MATLAB custom software (MathWorks, Inc. Natick, MA, USA) for post-processing behavioral data. We used the CoLBeRT system (54) to perform spatially-selective optogenetic manipulation for different motor circuit components. For optogenetic inhibition, we used a 561-nm solid-state laser with maximum intensity at 16 mW / mm^2^ (Arch) or a 473-nm solid-state laser at 0.76 mW / mm^2^ (GtACR2).

In the relevant figure legends, the *n* numbers, in most cases, denote the numbers of measurements on indicated numbers of different animals for the experiments, and represent both independent biological replication and technical replication.

### Optogenetic ablation

Optogenetic ablation was carried out using transgenic strains, in which mitochondrially targeted miniSOG (mini singlet oxygen generator) was specifically expressed in *C. elegans* neurons. Upon blue light illumination, mito-miniSOG caused rapid cell death in a cell-autonomous manner (34). To ablate AVB neurons [*Plgc*-55(B)::miniSOG or *Psra-11*::miniSOG], L2 / early L3 worms cultivated on OP50 were transplanted to an unseeded NGM plate, restricted within a 1.7 cm^2^ area via filter paper with a hole in the center and soaked with 100 μM CUCL_*2*_. Worms were illuminated with blue LED light (M470L3-C5, Thorlabs, Inc., USA) with an intensity of 80.2 mW / cm^2^, measured by a power meter (PM16-130, Thorlabs, Inc., USA). The temporal sequence was 0.5 / 1.5 s on / off pulses for 30 min. After illumination, worms were transplanted to newly OP50-seeded NGM plates with / without ATR for behavioral experiments. Mock ablations were performed via green light illumination using the same genotype under otherwise the same experimental conditions.

For selective B-type motor neuron ablation (*Pacr-5*::miniSOG), single L3 worms were transferred from the OP50-seeded NGM plates to 3% agarose (wt / vol) coated glass slides. The worms were covered by a cover glass to remain stationary. Spatially selective illumination patterns were generated by a digital micromirror device (DLI4130 0.7 XGA, Digital Light Innovations, TX, USA) to target individual neurons through a 20× objective mounted on a Nikon inverted microscope (Ti-U, Japan). Neurons were identified using mCherry fluorescence signals. We used a 473-nm blue laser with an intensity of 29 mW / mm^2^. The temporal sequence was 0.5 / 1.5 s on / off pulses for 15 min. After illumination, the worms were recovered by 2 μl of M9 buffer, and then transferred to OP50-seeded NGM plates with / without ATR for behavioral experiments.

## Acknowledgments

The authors thank Christopher Fang-Yen and Anthony Fouad for helpful discussion. This work was funded by the CAS Hundred Talents Plan and the National Science Foundation of China (NSFC-31471051 and NSFC-91632102 to Q. Wen) and the Canadian Institute of Health Research (CIHR-MOP93619, MOP123250 to M. Zhen).

## Supporting Information (SI)

### Linear model for undulatory wave propagation

We first developed a linear coupling model to simulate wave propagation along the body of *C. elegans*. In this model, the head motor circuit contains an oscillator that dominates the rhythmic motion of the whole body. Other segments cannot oscillate by themselves, but oscillate according to the proprioceptive input from the anterior segment. This simple phenomenological model does not simulate the detailed dynamics of individual neurons, but instead aims to elucidate several key properties of wave propagation based on a linear model.

The worm head motor circuit consists of a different group of motor neurons. Indeed, when all B-type motor neurons were optogenetically ablated, head rhythmic activity persisted during forward locomotion. Because neural mechanisms for generating head bending activity remain largely elusive, we, for simplicity, imposed sinusoidal head bending curvature for *k*_*1*_ at defined temporal frequency. Apart from the head, we divided the worm body into five segments (the number of segments was not essential for our theoretical argument), each with different dynamics, as described in Equation 1. Equation 1 also arises naturally from a biologically more realistic model, where dorsal and ventral activities are treated separately. In this case, we have:

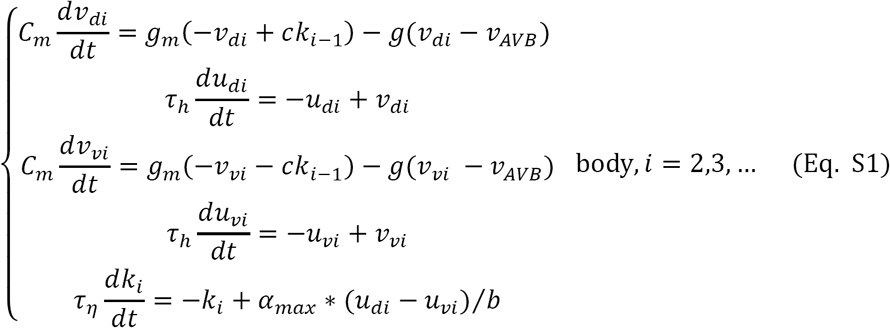

Subscripts *d* and *v* denote dorsal and ventral sides, respectively. If we subtract “dorsal” equations from “ventral” equations, Equation 1 will be derived, provided we consider that dorsal and ventral motor activities are anti-phased, namely *v*_**d**_ = -*v*_**i**_ and *v*_**di**_ = -*v*_**vi**_. Simulation using Eq. S1 did not show any essential difference compared with simulation using Equation 1.

We set AVB membrane potential at a constant depolarized value in our model because AVB has gap junction coupling with all B-type motor neurons. During forward locomotion, B-type neurons are at distinct oscillatory phase: some are depolarized and some are hyperpolarized. The net electrical currents AVB received are cancelled out and remain roughly constant. This assumption is consistent with calcium imaging data of AVB, which exhibited elevated yet non-oscillatory activity during forward locomotion.

The linear model makes two predictions that can be tested experimentally. First, abolishing head motor activity should eliminate undulatory wave propagation. However, when the head and body were decoupled in our experiments, we observed intrinsic high frequency oscillation emerging from the mid-body. Second, while simulation suggested AVB-B gap junction coupling would deteriorate the propagation of the bending amplitude (Fig. 2*D*), our experimental data revealed the opposite. This result was independent of the value of *v*_*AVB*_.

Parameters used in simulation are:

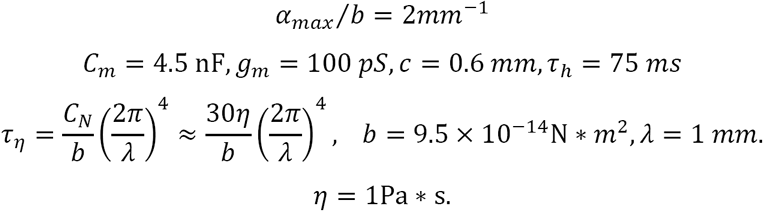

### Analytical solution to linear model

To gain a deeper insight into the bending amplitude decay during wave propagation (Fig. 2*D*), we developed an analytical understanding for the linear coupling model by considering the following continuous model:

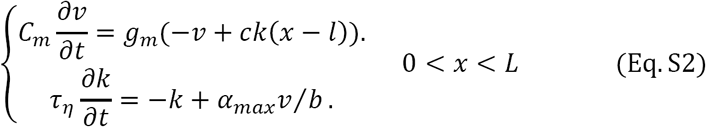

In the first equation, *ck*(*x* - *l*) describes how proprioceptive couplings drive the change in neural activity in motor neurons. *l* is the characteristic length of proprioceptive coupling from an anterior body region, and *L* is the length of the worm. The second equation relates the change of body shape with neural activity. For simplicity, we dropped the equation that relates neural activity and muscle torque in Equation 1. The delay time between neural activity and muscle torque, characterized by τ_**h**_, was absorbed by the time constant τ_**η**_ in Eq. S2. Because proprioceptive coupling is local, we impose *l* ≪ *L*.

We assert a harmonic motion to describe the motor activity of the head. Thus, *v*(*x*) and *k*(*x*) can be written as:

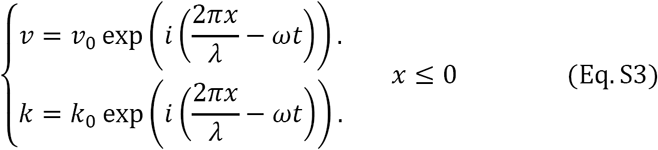

The neural activity and curvature are not in-phase. Thus, the phase factors are absorbed into coefficients *v*_*0*_ and *k*_*0*_, which are all complex numbers.

The self-consistent solution for the motor activity along the body is sinusoidal, namely

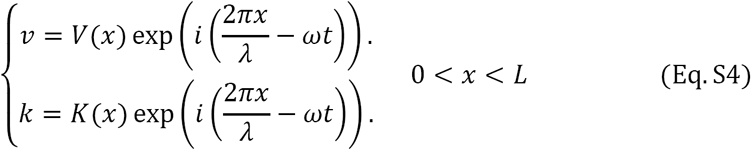

Substituting Equation 11 into Equation 9 and defining τ_**m**_ = *C*_**m**_ / *g*_**m**_, we have

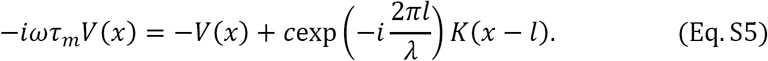

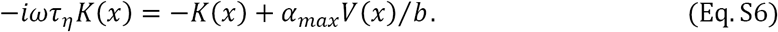

Replacing *K*(*x* - *l*) with *K*(*x*) - *K*′ (*x*)*l* + *K*″ (*x*)*l*_*2*_ / 2, and using *l* ≪ λ, we find

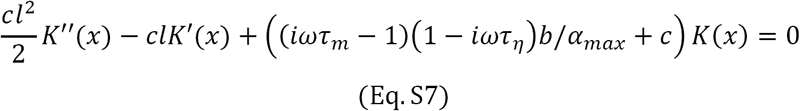

The solution is

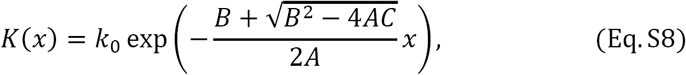

where

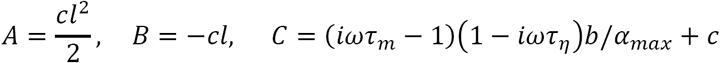

Thus, the decaying length constant satisfies

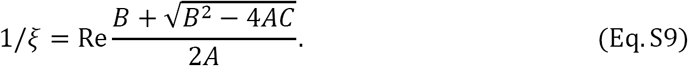

Expanding it to power series of *ω* and only preserving lower order terms, we have

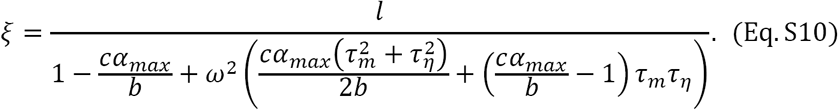

The time constant τ_*η*_ increases with viscosity of medium η, and the decay length constant ξ decreases with η.

To the leading order, the decay length constant can be approximated as

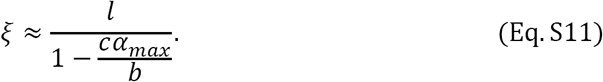

Note that Eq.S11 is consistent with the discrete model with nearest neighbor coupling, at least to the leading order. In the discrete model of nearest neighbor coupling, the curvatures follow the relationship κ_**i**_ ≈*cα*_**max**_/ bκ_**i-1**_ (31). So the decaying rate should be 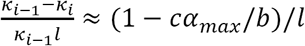, which is identical to the result from the continuous model.

When we incorporate electrical couplings between AVB interneurons and B-type neurons into this model, the equations become

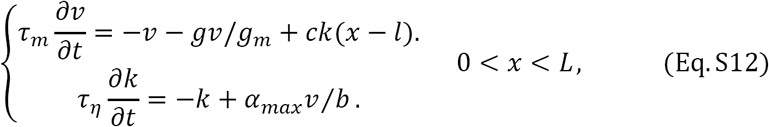

where *g* is the gap junction conductance. In this case, the decay length constant, to the leading order, reduces to:

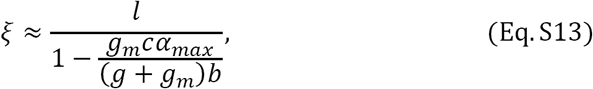

which further deteriorates the propagation of the bending amplitude. Note that although the effective τ_**m**_ is also smaller, 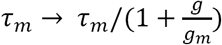, its contribution is mostly to the second order and does not change the conclusion. This prediction is inconsistent with our experimental observations.

### Nonlinear model

The discrepancy between our theoretical and experimental observations forced us to develop a modified nonlinear model. For the head oscillator, we still imposed dorsal-ventral anti-correlated bending activities at a defined temporal frequency. For all other segments, our model incorporated the experimental observations that B-type motor neurons generate intrinsic oscillations when the head and body are decoupled and when AVB interneurons and B-type motor neurons are electrically coupled (*g* > 0). The intrinsic membrane properties in the B-type motor neurons remain unknown. For simplicity, we developed a phenomenological model based on active Ca^2+^ and K^+^ conductances (41).

The motor activity in a given body segment is now governed by the following equations:

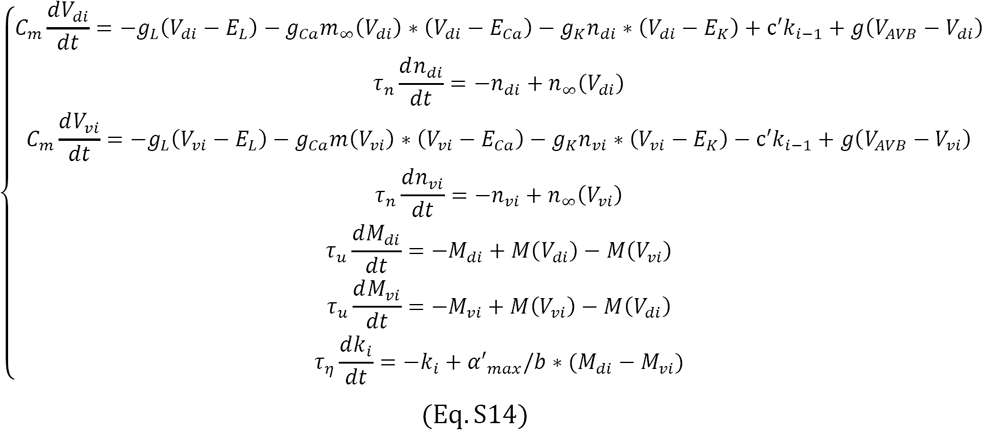

In Eq. S14, *m*, *n* are voltage-dependent gating variables; *m*_*∞*_(*V*), *n*_*∞*_(*V*) are steady-state activation functions. We assume that the time constant for the gating variable *m* is small and *m* follows instantaneously with the voltage. *M*(*V*) relates cell membrane potential and muscle torque. We assume they all have sigmoidal functional forms:

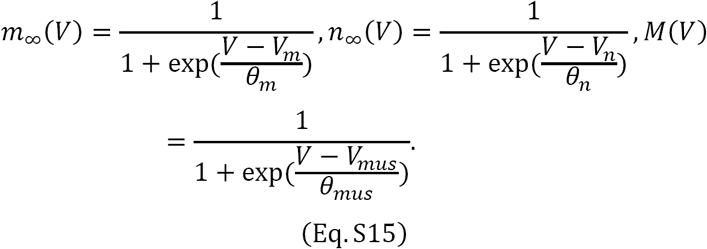

*g*(*V*_*AVB*_ - *V*) is the current flowing through the gap junction between AVB interneurons and B-type neurons, where *V*_*AVB*_ is the membrane potential of the AVB interneurons, held at a constant value.

The terms -*M*(V_**v**_) and -*M*(*V*_**d**_) in Eq. S14 reflect contralateral inhibition from the D-type GABAergic motor neurons: dorsal D neurons can be activated by ventral B neurons and thereby inhibit dorsal muscle activity, and vice versa.

We simulated six segments (including the head), similar to that in the linear coupling model. The segment number was roughly in accordance with the number of DB-type motor neurons. The parameter values are listed below.

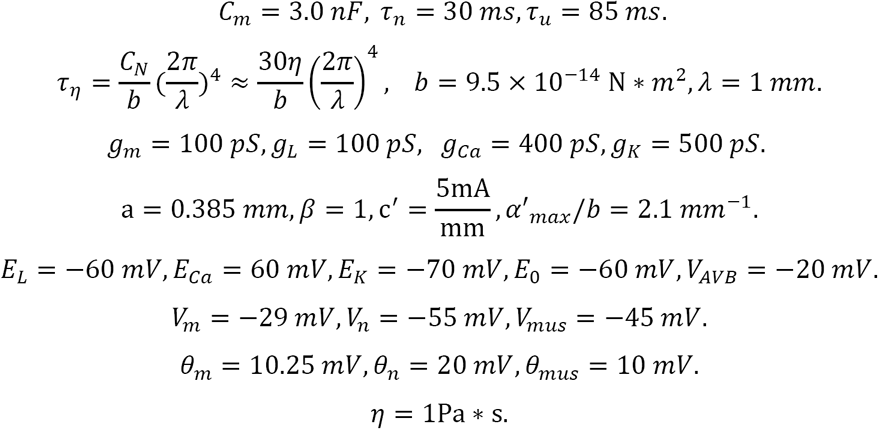

Comparing the case *g* = 0 and *g* = 100 *pS*, we found that gap junction coupling between AVB interneurons and B-type neurons could equalize the bending amplitude (Fig. 5*C*). High frequency oscillation within middle segments could also be generated when the head and body were decoupled (Fig. 5*A*).

### Videos

**Video 1** | An *unc-7* mutant animal swam freely in solution at ∽ 1 Pa·s viscosity. Despite anterior body undulation, the bending amplitude near the tail significantly diminished.

**Video 2** | Optogenetic inhibition of the anterior region of a freely swimming wild type worm induced high frequency undulation in the posterior body region.

**Video 3** | A worm with UNC-7 expression restored in AVB and other neurons (P*sra-11*::UNC-7) swam in solution at ∽ 1 Pa·s viscosity. Both bending amplitude and high-frequency undulation in the posterior body region were restored.

**Video 4** | Optogenetic inhibition of the mid-body B-type motor neurons in a wild type worm resulted in whole body paralysis.

**Video 5** | Optogenetic inhibition of the mid-body B-type motor neurons in an *unc-9* mutant worm led to paralysis in the posterior region, whereas the anterior body undulation persisted.

**Video 6** | Over-expression of UNC-9 innexin in B-type motor neurons (P*acr-5*::UNC-9) caused whole body paralysis. Worms were incapable of propagating body undulations along their bodies.

**Video 7** | Optogenetic activation of AVB premotor interneurons (P*sra-11*::Chrimson) could trigger forward locomotion from the pause state.

**Video 8** | Optogenetic inhibition of AVB premotor interneurons (P*sra-11*::GtACR2) could abolish forward locomotion.

Note that all videos were played at 0.5x speed.

**Fig. S1.**
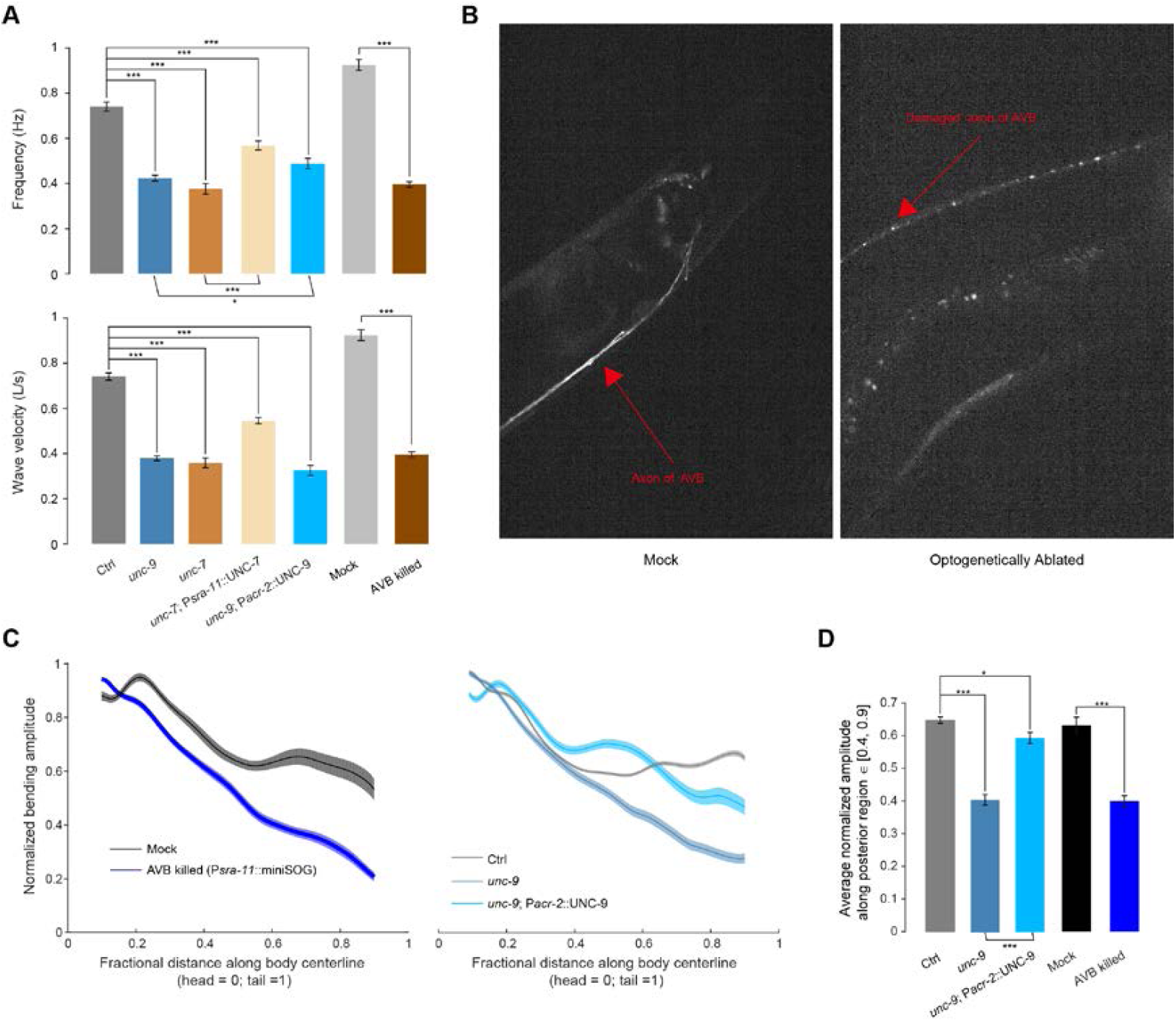
AVB-B electrical couplings facilitate undulatory wave propagation during forward locomotion. (*A*) Locomotion kinematics of tested strains. Upper panel shows the comparison of undulation frequency in viscous medium (∽ 1 Pa·s). Lower panel shows the comparison of wave velocity, calculated in worm body length per second (*L* / *s*). Error bars show s.e.m. *** *p* < 0.0001, * *p* = 0.01, two-sample *t-*test with Bonferroni correction. Ctrl [N2; P*acr-5*::Arch; P*unc-4*::TWK-18(gf)], *n* = 107 measurements, 13 worms; *unc-7* mutant, [*unc-7*(*hp121*); P*acr-5::*Arch; P*unc-4*::TWK-18(gf)], *n* = 99 measurements, 22 worms; *unc-9* mutant [*unc-9*(*fc16*); P*acr-5*::Arch; P*unc-4::*TWK-18(gf)], *n* = 144 measurements, 17 worms; AVB ablated worm [P*glc-55(*B)::tom20::miniSOG::UrSL::wCherry], *n* = 80 measurements, 13 worms; Mock, *n* = 96 measurements, 12 worms. (*B*) Comparison of AVB axon fluorescence signals in mock and AVB-ablated worms. Arrows indicate intact and damaged axons. (*C*) Left: Comparison of normalized bending amplitude along the worm body in mock and AVB-ablated worms. We used a different transgenic strain (P*sra-11*::miniSOG) to drive the expression of miniSOG in AVB neurons. Right: Comparison of normalized bending amplitude along the worm body in CTRL, *unc-9*, and UNC-9 rescue in B-type and A-type motor neurons [*unc-9*(*fc16*); P*acr-2*::UNC-9; P*acr-5*::arch]. Rescue of UNC-9 in B-type and A-type motor neurons largely restored the posterior bending amplitude. Shaded regions show s.e.m. (*D*) Bending amplitude averaged over the posterior region ∈ [40, 90] of the worm body. \*\*\**p* < 0.0001, \**p* = 0.003, two-sample *t*-test with Bonferroni correction. Error bars represent s.e.m. Ctrl, *n* = 105 measurements, 13 worms; *unc-9* mutant, *n* = 93 measurements, 17 worms; UNC-9 rescue in motor neurons, *n* = 64 measurements, 9 worms. Mock, *n* = 28 measurements, 5 worms; AVB-ablated worms (P*sra-11*::tomm20::miniSOG) *n* = 100 measurements, 10 worms.

**Fig. S2.**
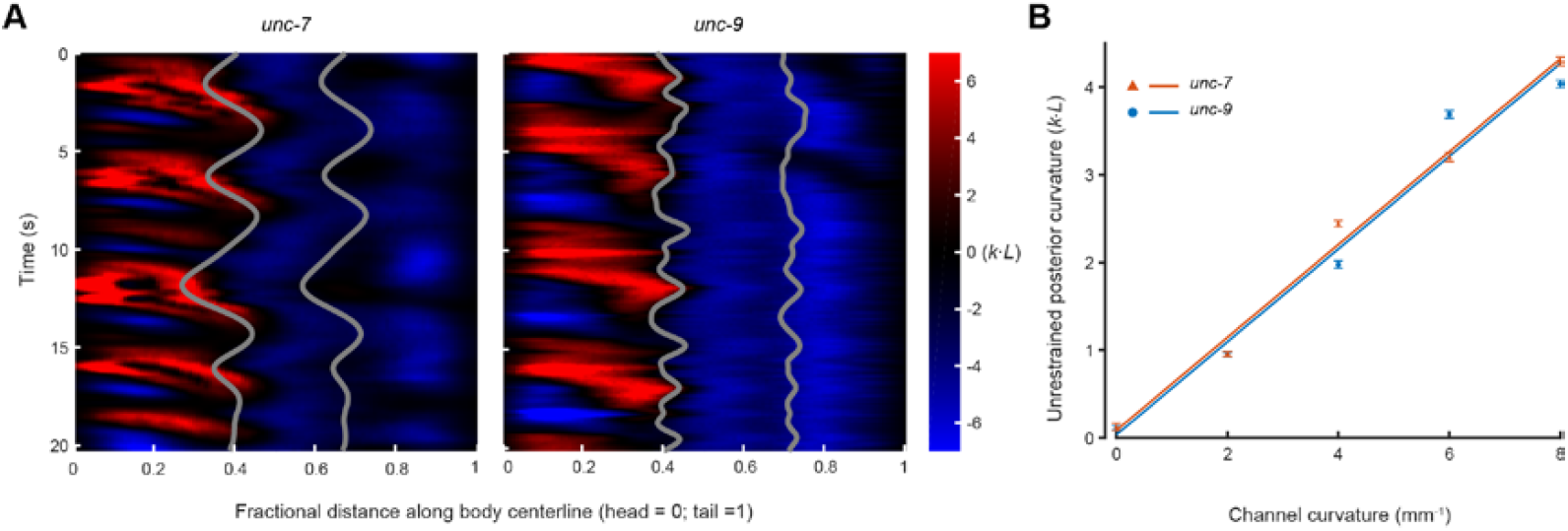
AVB-B electrical couplings are not necessary for transducing proprioceptive signals. (*A*) Representative kymographs showing an *unc-7* mutant [*unc-7*(hp121); P*unc-4::*TWK-18(gf)] and an *unc-9* mutant worm [*unc-9*(fc16); P*unc-4::*TWK-18(gf)], whose mid-bodies were restrained in a microfluidic channel with defined curvature (6 mm^-1^). Gray lines indicate anterior and posterior boundaries of the channel. (*B*) Unrestrained posterior body curvature as a function of channel curvature. Each data point: *n* = 5 worms, error bars represent s.e.m. Blue and red lines are linear fit (*R*^2^ > 0.97).

**Fig. S3.**
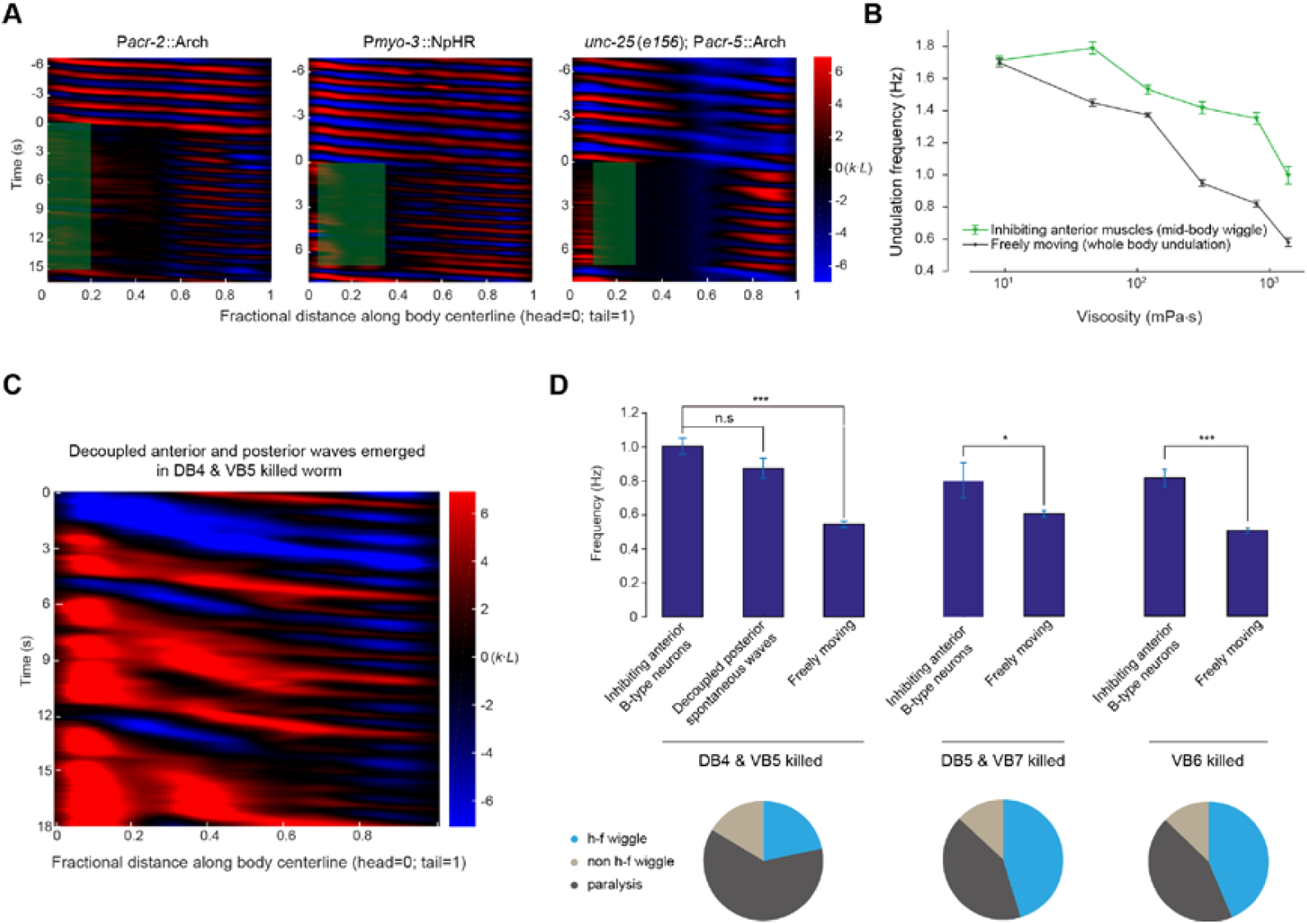
Mid-body B motor neurons intrinsically generate rhythmic activity. (*A*) Representative kymographs demonstrate that fast mid / posterior body undulatory waves could be induced by inhibiting anterior A-&B-type motor neurons (P*acr-2*::Arch, 11 out of 25 measurements) by inhibiting anterior muscle cells (P*myo-3*::NpHR, 52 out of 87 measurements) or by inhibiting anterior B-type motor neurons in GABA neurotransmitter-deficient worms [*unc-25(e156)*; P*acr-5*::Arch, 33 out of 45 measurements]. (*B*) Undulation frequency with medium viscosity. Black, normal swimming worm; green, mid-body undulation when the head and body were decoupled by inhibiting anterior muscle cells. Each data point represents mean and s.e.m from n ≥ 4 worms. (*C*) Representative kymographs of a freely swimming worm in which DB4 & VB5 motor neurons were optogenetically ablated (P*acr-5*::tomm20::miniSOG). Decoupled spontaneous waves emerged from anterior and posterior body regions. (*D*) Up, summary of undulation frequencies after selectively ablating a subset of B-type motor neurons in the mid-body. Three behavioral outcomes were compared. (1) Fast mid-body undulation induced by optogenetically inhibiting anterior B-type motor neurons. (2) Spontaneous posterior bending waves in DB4&VB5-ablated worms. (3) Freely swimming worms. Bottom, pie chart summarizes worm locomotor states across trials during optogenetic inhibition of anterior B-type motor neurons. State definitions can be found in the main text. Error bars show s.e.m., *** *p* < 0.0001, * *p* = 0.01, *n.s. p* = 0.15, two-sample t-test. DB4 & VB5 killed n = 11 worms, DB5 & VB7 killed n = 8 worms, VB6 killed n = 9 worms.

**Fig. S4.**
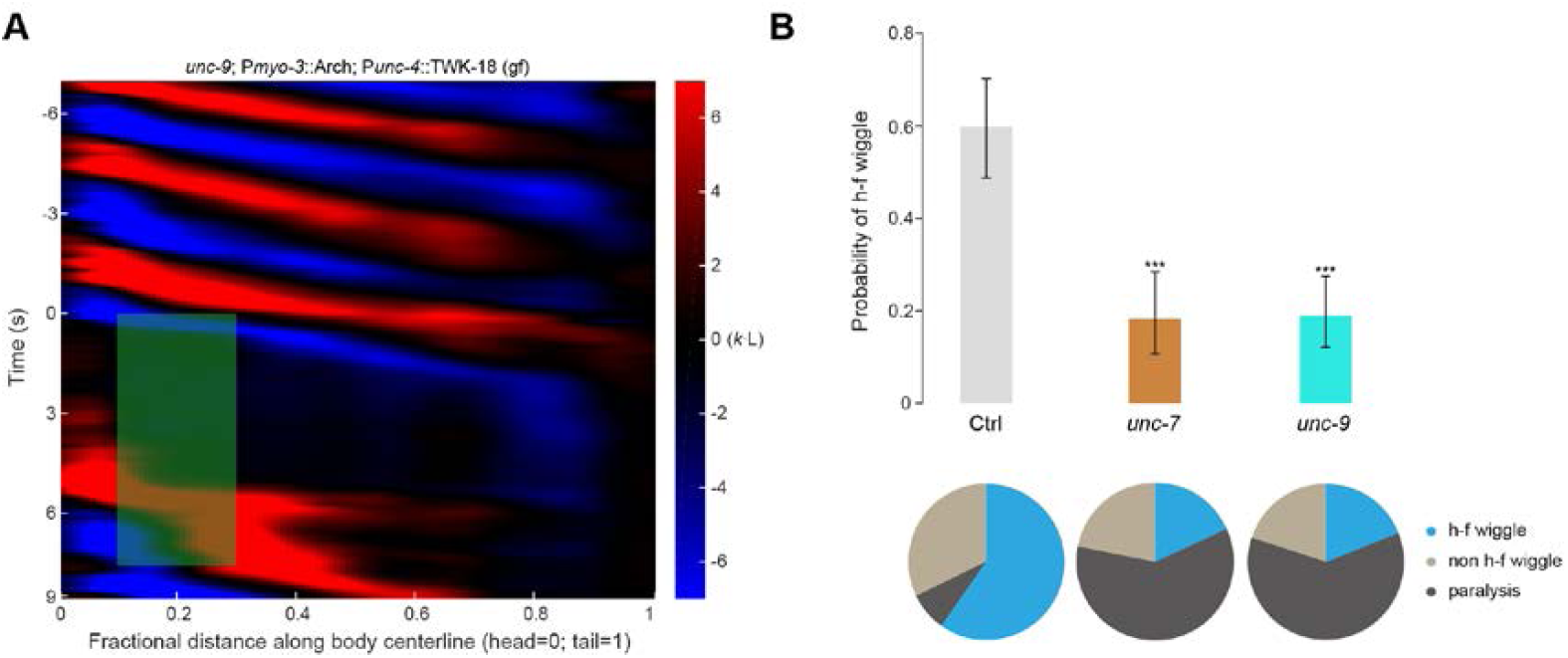
AVB-B electrical couplings drive a bifurcation in B-type neuron dynamics. (*A*) Representative curvature kymographs of head-body-decoupled experiments in AVB-B gap junction-deficient *unc-9* mutants [*unc-9*; P*myo-3*::Arch; P*unc-4*::TWK-18(gf)]. Anterior muscle cells were optogenetically inhibited to abolish the proprioceptive signals arising from head bending activity. (*B*) Probability for inducing high frequency mid-body undulation when the head and body were decoupled by inhibiting anterior muscle cells. Pie chart summarizes worm locomotor states across trials. Error bars indicate 95% binomial proportion confidence interval. *** *p* < 0.0001, Chi-Square test. Ctrl [P*myo*-3::NpHR, P*unc-4::*TWK-18(gf)], *n* = 87 measurements, 10 worms. *unc-7* mutant [*unc-7(hp121);* P*myo*-3::Arch; P*unc-4::*TWK-18(gf)], *n* = 82 measurements, 18 worms. *unc-9* mutant [*unc-9(fc16);* P*myo*-3::Arch; P*unc-4::*TWK-18(gf)], *n* = 111 measurements, 21 worms.

**Fig. S5.**
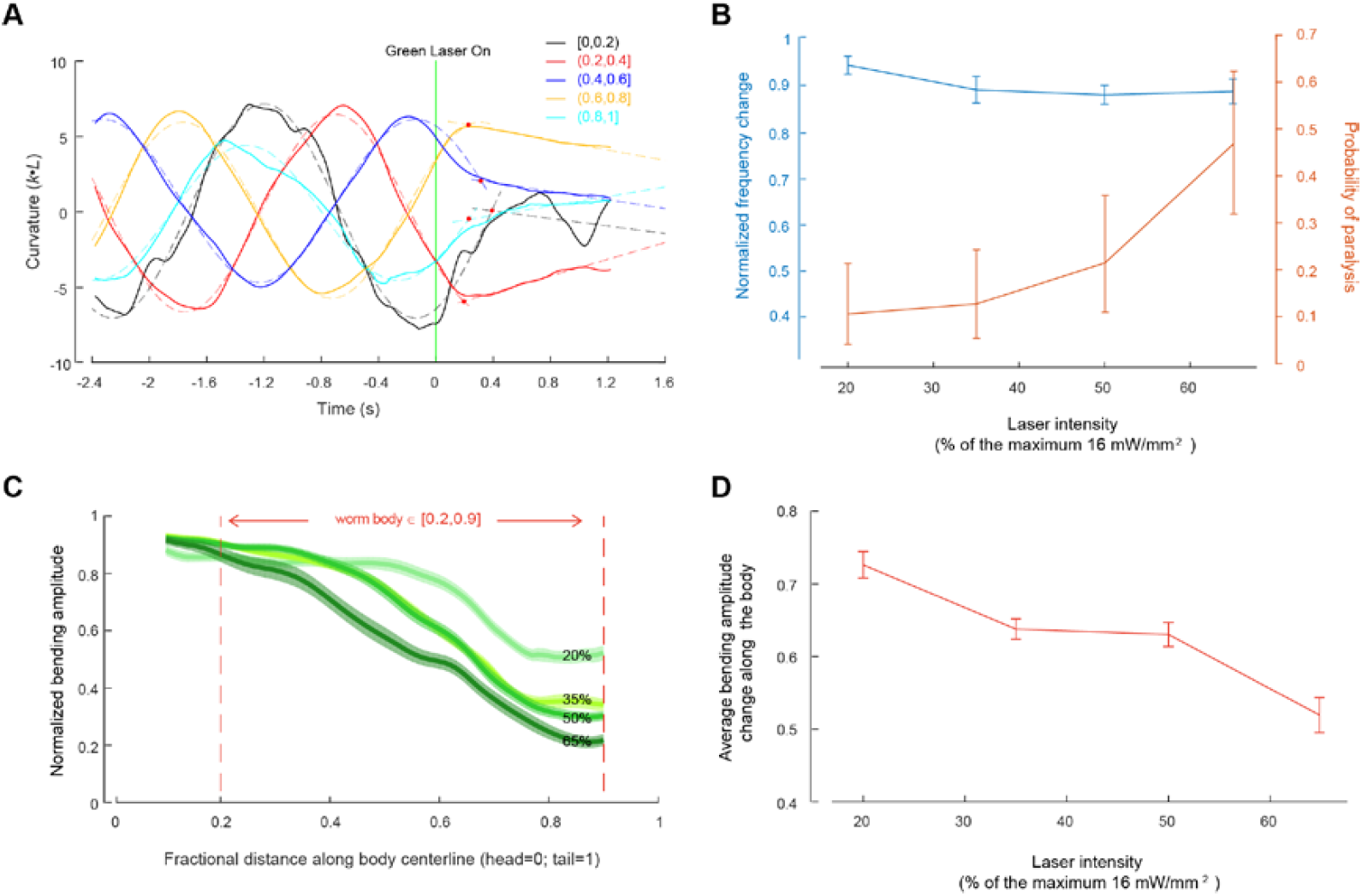
Electrical couplings between motor neurons allow for rapid and reciprocal interactions between head and body motor activities. (*A*) Estimation of reaction time when mid-body B-type motor neurons were optogenetically inhibited. First, we fit the regional curvature with a sinusoidal function right before the green laser was turned on. Second, we performed a linear fit of regional curvature during the optogenetic inhibition. The crossover point (red dot) represents the wiggle-to-paralysis transition for each segment. Curvatures were averaged over a defined color-coded region, represented by a fractional distance along the worm body coordinates (head = 0, tail = 1). (*B*) Dose-dependent effect on the undulation frequency and efficacy for inducing whole-body paralysis when B-type motor neurons within a posterior body region (∈ [0.7, 0.9]) were optogenetically inhibited. Error bars (blue), s.e.m; error bars (orange), 95% binomial proportion confidence interval. *n* ≥ 5 worms for each data point. (*C*) Dose dependent effect on the bending amplitude along the worm body when B-type motor neurons within a posterior body region (∈ [0.7, 0.9]) were optogenetically inhibited. Shaded regions are s.e.m. (*D*) Average bending amplitude change along the body ∈ [0.2, 0.9]. Error bars are s.e.m. Maximum laser intensity was 16 mW / mm^2^.

**Table S1.** Strain information

| Strain Name | Experiment | Genotype | Construct (plasmid no.); injection concentrations |
| --- | --- | --- | --- |
| WEN0315 | B motor neuron inhibition in WT | *wenIs0001*;<br>*wenEx0315* | *Pacr-5::Arch::UrSL::wCherry* (pJH2918); *Podr-1::gfp* (wenIs0001)<br>*Punc-4::TWK::18(gf)::UrSL::wCherry* (pJH2108) 30 ng/μL; *Plin-14::gfp* 20 ng/μL; *Pmyo-3::wCherry* (pJH1774) 2.5 ng/μL (wenEx0315) |
| WEN0317 | B motor neuron inhibition in *unc-7* mutants; Microfluidic experiments | *wenIs0001*;<br>*wenEx0317*;<br>*unc-7(hp121)* | *Pacr-5::Arch::UrSL::wCherry* (pJH2918); *Podr-1::gfp* (WenIs0001)<br>*Punc-4::TWK::18(gf)::UrSL::wCherry* (pJH2108) 30 ng/μL; *Plin-14::gfp* 20 ng/μL; *Pmyo-3::wCherry* (pJH1774) 2.5 ng/μL (wenEx0317) |
| WEN0304 | B motor neuron inhibition in *unc-9* mutants; Microfluidic experiments | *wenEx0304*;<br>*hpEx2088*;<br>*unc-9(fc16)* | *Pacr-5::Arch::UrSL::wCherry* (pJH2918) 30 ng/μL; *Plin-14::gfp* 20 ng/μL (Wen Ex0304)<br>*Punc-4::TWK::18(gf)::UrSL::wCherry* (pJH2108) ; *Podr-1::gfp* (hpEx2088) |
| ZM7297 | AVB ablation | *hpls331*; | *Plgc-55(B)::tomm-20::miniSOG::UrSL::wCherry* (pJH2890); *lin-15* |
| WEN0318 | B motor neuron inhibition upon AVB ablation; Microfluidic experiments | *wenIs0001*;<br>*hpls331*; | *Pacr-5::Arch::UrSL::wCherry* (pJH2918); *Podr-1::gfp* (wenIs0001)<br>*Plgc-55(B)::tomm-20::miniSOG::UrSL::wCherry* (pJH2890) (*hpls331*) |
| WEN0335 | B motor neuron inhibition in *unc-7* mutants, with UNC-7 expression restored in AVB | *wenEx0335*;<br>*wenIs0001*;<br>*unc-7(hp121)* | *Punc-4::TWK::18(gf)::UrSL::wCherry* (pJH2108) 30 ng/μL; *Psra-11::UNC-7(cDNA)::wCherry* (pJH2037) 20 ng/μL; *Plin-14::gfp* 20ng/μL; *Pmyo-3::wCherry* 2.5 ng/μL (wenEx0335)<br>*Pacr-5::Arch::UrSL::wCherry* (pJH2918); *Podr-1::gfp* (wenIs0001) |
| WEN0314 | B motor neuron ablation; Microfluidic experiments | *wenIs0001*;<br>*hpls374*; | *Pacr-5::Arch::UrSL::wCherry* (pJH2918); *Podr-1::gfp* (*hpls374*)<br>*Pacr-5::tomm-20::miniSOG::UrSL::wCherry* (pJH2842) (wenIs0001) |
| WEN0001 | B motor neuron inhibition | *wenIs0001* | *Pacr-5::Arch::UrSL::wCherry* (pJH2918); *Podr-1::gfp* (wenIs0001) |
| WEN0320 | B motor neuron inhibition upon AVB ablation | *wenIs0001*;<br>*wenEx0320* | *Pacr-5::Arch::UrSL::wCherry* (pJH2918); *Podr-1::gfp* (wenIs0001)<br>*Psra-11::tomm-20::miniSOG::UrSL::wCherry* (quan0065) 50 ng/μL; *Pmyo-3::wCherry* 2.5 ng/μL (wenEx0320) |
| ZM9275 | A & B motor neuron inhibition | *hpls615* | *Pacr-2(s)::Arch::wCherry* (pJH3717) (*hpls615*) |
| ZX444.6 | Muscle inhibition | *zxEx29*;<br>*lin-15(n765ts)* | *Pmyo-3::NpHR::ECFP*; *lin-15+* (*zxEx29*) |
| WEN0307 | B motor neuron inhibition in GABA deficient mutants | *wenIs0001*;<br>*unc-25(e156)* | *Pacr-5::Arch::UrSL::wCherry* (pJH2918); *Podr-1::gfp* (wenIs0001) |
| ZM7796 | B motor neuron ablation | *hpls374* | *Pacr-5::tomm-20::miniSOG::UrSL::wCherry* (pJH2842) (*hpls374*) |
| WEN0372 | AVB activation | *wenEx0372* | *Psra-11::Chrimson::wCherry* (quan0164) 30 ng/μl; *Plin-24::GFP* (*wenEx0372*) |
| WEN0387 | AVB inhibition | *wenEX0387* | *Psra-11::GtACR2::wCherry* (quan0235) 30 ng/μl; *Plin-24::GFP* (*wenEX0387*) |
| WEN0363 | Muscle inhibition in *unc-7* mutants | *wenEX0363*; *unc-7(hp121)* | *Pmyo-3::Arch::UrSL::wCherry* (quan0124) 30 ng/μl; *Punc-4::TWK-18(gf)::UrSL::wCherry* (pJH2108) 30 ng/μl (*wenEx0363*) |
| WEN0323 | Muscle inhibition in *unc-9* mutants | *wenEx0323*; *unc-9(fc16)* | *Pmyo-3::Arch::UrSL::wCherry* (quan0124) 30 ng/μl; *Punc-4::TWK-18(gf)::UrSL::wCherry* (pJH2108) 30 ng/μl (*wenEx0323*) |
| WEN0337 | *unc-9* mutants with restored expression in A and B motor neurons | *wenEx0337*; *unc-9(fc16)* | *Pacr-2::UNC-9::wCherry* (pJH1897) 2 ng/μl; *Pacr-5::Arch::UrSL::wCherry* (pJH2918) 30 ng/μl (*wenEx0337*) |
| WEN0366 | Overexpression of UNC-9 in B motor neurons | *wenEx0366* | *Pacr-5::Arch::UrSL::wCherry* (pJH2918) 30 ng/μl; *Pacr-5::UNC-9* (pJH2140) 30 ng/μl (*wenEx0366*) |

